# Programmable Imaging of Locus-Specific DNA Repair Dynamics Reveals Chromatin-Controlled Therapy Resistance in Cancer

**DOI:** 10.64898/2026.11.12.759262

**Authors:** Linshan Zhu, Abdula Monther, Anthony Mouchawar, Mishel Tsoy, Xi Yu, Ziliang Huang, Qin Peng, Yunjia Qu, Yiqian Wu, Longwei Liu, Yingxiao Wang

## Abstract

DNA double-strand breaks (DSBs) pose a significant threat to genomic integrity, and their efficient repair is critical for cell survival. However, the spatial and temporal aspects of the repair process at specific genomic loci remain a mystery. To address this gap, we developed a novel live-cell imaging platform called <u>G</u>enomic <u>L</u>ocus Imaging and <u>D</u>SB <u>R</u>epair Monitoring by <u>E</u>lastin-like Proteins (GLIDRE). This system leverages a CRISPR-guided elastin-like protein (ELP) droplet approach to specifically label non-repetitive genomic loci and enables real-time analysis of repair-associated dynamics at the same site. Using GLIDRE, we demonstrate that DSB repair-associated dynamics are significantly faster in open chromatin regions compared to compacted regions, irrespective of local transcriptional activity, supporting a model in which chromatin accessibility defines the baseline response whereas transcriptional activity acts in a context-dependent manner. In a model of diffuse intrinsic pontine glioma (DIPG), the H3.3K27M mutation reshapes chromatin accessibility and repair dynamics in a locus-specific manner. At the tumor suppressor locus CDKN2A, H3.3K27M mutation caused a more repressive chromatin state and delayed repair-associated dynamics, which was associated with resistance to CAR T-cell therapy. Pretreatment with the histone deacetylase inhibitor suberoylanilide hydroxamic acid (SAHA) restores repair dynamics at CDKN2A and sensitizes H3.3K27M tumor spheroids to CAR T-cell therapy. Together, our findings reveal chromatin accessibility as a key determinant of DSB repair dynamics and support a mechanistic link and actionable connection between chromatin state, DNA repair, and immunotherapy response.

## Introduction

DNA damage is one of the most critical challenges faced by cells, as it can lead to genomic instability, cell death, and various pathological conditions, including cancer ^1^. Among different types of DNA damage, double-strand breaks (DSBs) are particularly hazardous due to their potential to cause chromosomal rearrangements and mutations if not properly repaired ^2^. The repair of DNA damage involves complex mechanisms that must balance both speed and accuracy to maintain genomic integrity ^3,4^. Lesions must be repaired fast enough for cells to resume normal function. Cells that fail to repair lesions in a timely manner usually undergo apoptosis, and cells with hyperactive repair capacity, including some cancer cells, can acquire resistance to genotoxic treatment ^5,6^.

Traditional methods for studying DNA repair have largely relied on fixed-cell imaging techniques, particularly through γH2AX immunofluorescence ^7^. Quantitative live-cell imaging approaches have provided important insight into the dynamics of molecular recruitment and damage responses in living cells ^8^. However, these methods generally do not provide locus-specific information at endogenous non-repetitive genomic sites, and the spatiotemporal dynamics of repair at defined genomic loci therefore remain difficult to resolve in living cells ^9,10^. This limitation has driven the development of new tools for live-cell imaging of DNA loci. For instance, CRISPR-based systems have been engineered to label specific genomic loci in living cells, typically using tandem MS2 loops on gRNAs or SunTag amplification on dCas9 to enhance target signals ^11–16^. However, low resolution on non-repetitive sequences due to insufficient signal amplification remains a major challenge for the visualization and tracking of vital genes, many of which do not contain repetitive target sequences.

Liquid-liquid phase separation (LLPS) has emerged as a powerful mechanism for locally concentrating biomolecules through self-assembly, thereby amplifying molecular signals within cells and has become an important conceptual framework across physics, chemistry, and biology ^17^. Recent studies, including our own, have exploited foldon-and HP1a-based LLPS systems to label non-repetitive genomic loci in living cells ^18,19^. These findings suggest that engineered phase-separating scaffolds can overcome the weak signal limitations of conventional locus-labeling strategies. Elastin-like polypeptides (ELPs), synthetic polymers composed of repetitive [VPGXG] motifs derived from elastin, undergo tunable phase transitions depending on sequence composition, concentration, and environmental conditions ^20–22^. Because of their biocompatibility, modularity, and well-characterized phase behavior, ELPs have been widely used in tissue engineering and drug delivery ^23,24^. We therefore reasoned that ELPs could serve as an attractive LLPS scaffold for live-cell imaging of non-repetitive genomic loci.

Brain tumors remain one of the most challenging cancers to treat, particularly in pediatric patients. Diffuse intrinsic pontine glioma (DIPG), characterized by the presence of H3.3K27M mutations, represents one of the most aggressive and lethal forms of pediatric brain cancer ^25^. The resistance of these tumors to conventional therapies such as chemotherapy and radiotherapy has prompted intensive research into understanding their molecular mechanisms of survival and developing more effective treatment strategies ^26^. Recent advances in cancer immunotherapy, particularly chimeric antigen receptor (CAR) T-cell therapy, have shown promising results in various malignancies ^27,28^. However, similar to conventional therapies, H3.3K27M-mutant DIPG often exhibits limited or non-durable responses to CAR T-cell therapy, and the mechanisms underlying this resistance remain incompletely understood ^27–29^. Resistance to both radiotherapy and CAR T therapy may involve overlapping tumor-intrinsic survival mechanisms. In H3K27M DIPG, radiotherapy response is tightly linked to DNA damage and repair capacity, and emerging studies suggest that DNA damage response pathways can also influence CAR T efficacy and may represent rational combination targets ^30–33^.

In this study, we introduce the GLIDRE (<u>G</u>enomic <u>L</u>ocus Imaging and <u>D</u>SB <u>R</u>epair Monitoring by <u>E</u>lastin-like Proteins) system, a live-cell imaging platform for the concurrent imaging of genomic loci and monitoring of DSB repair dynamics. Compared with existing live-cell locus imaging approaches, GLIDRE provides three technical advances. First, it enables robust visualization of endogenous non-repetitive genomic loci through CRISPR-guided, phase-separation-mediated signal amplification. Second, it integrates locus labeling with controlled local perturbation and simultaneous readout by combining orthogonal Cas12a-based DSB induction with live imaging of repair-associated chromatin responses at the same genomic region. Third, it enables time-resolved kinetic measurement of locus-specific damage responses in living cells, allowing dynamic comparison of chromatin-state-dependent behaviors across defined genomic coordinates. Using GLIDRE, we investigated the impact of transcriptional activity and chromatin accessibility on the DNA repair process and found that chromatin accessibility shows a stronger association with repair-associated dynamics than local transcriptional activity. Furthermore, we applied this platform to examine the mechanisms underlying the DNA repair dynamics in an H3.3K27M tumor model relevant to therapy resistance. Intriguingly, our results indicate that H3.3K27M reshapes chromatin accessibility and DNA repair dynamics in a locus-specific manner under genotoxic stress. At the tumor suppressor locus CDKN2A, H3.3K27M is associated with a more repressive chromatin state and delayed repair-associated dynamics, suggesting a potential mechanism that may contribute to the therapeutic resistance observed in these tumors. Importantly, we further show that pretreatment with suberoylanilide hydroxamic acid (SAHA), a histone deacetylase (HDAC) inhibitor, restores repair dynamics at CDKN2A and enhances sensitivity to CAR T-cell therapy, supporting chromatin-modulating strategies as a potential approach to overcome resistance in H3.3K27M-mutant tumors.

## Results

### 1. Engineering a CRISPR-guided ELP droplet system for genomic loci labeling in live cells

Elastin-like polypeptides (ELPs), known for their phase transition behavior, biocompatibility, and high tunability, have been extensively utilized in tissue engineering and drug delivery ^23,24^. To adapt ELPs for intracellular locus imaging, we selected the [VPGVG]_n_ variant (ELP-Vn), since its phase transition temperature (37–45°C) aligns well within physiological conditions ^34,35^ (**Figure 1a**). Codon-optimized variants of ELP-Vn (ELP-V10, V20, V30, V45, and V60) were constructed for phase transition test in HEK293T cells ^36^ (**Supplementary Figure S1a**). GFP was fused to the N-terminal of these ELP variants to visualize their behavior in live cells under the microscope ^37^ (**Figure 1b**). We found that variants with fewer than 30 repeats remained soluble, while GFP-ELP-V60 exhibited constant intracellular droplets at 37°C. GFP-ELP-V45-transfected cells displayed heterogeneous distribution, with some cells showing droplets, and others a diffuse GFP pattern (**Figure 1b and Supplementary Figure S1b**). We quantified droplet formation by measuring the total projected droplet area in each cell and plotted this against the corresponding GFP fluorescence intensity, which reflects the relative intracellular concentration of GFP-ELP-V45. Cells expressing low levels of GFP-ELP remained diffuse with minimal droplet area, while cells with progressively higher GFP levels displayed an increase in droplet area. This concentration–response trend demonstrates that intracellular ELP phase transition is concentration dependent (**Figure 1c**). To further validate the property of the droplets, we treated the cells with 2.5% 1,6-hexanediol (1,6-HD), a chemical compound commonly used to disrupt the liquid-liquid phase separation (LLPS) ^38,39^. Consistent with published data, the droplets completely dissolved after 1,6-HD treatment, verifying the LLPS property of ELPs (**Supplementary Figure S2a,b**).

**Figure 1.**
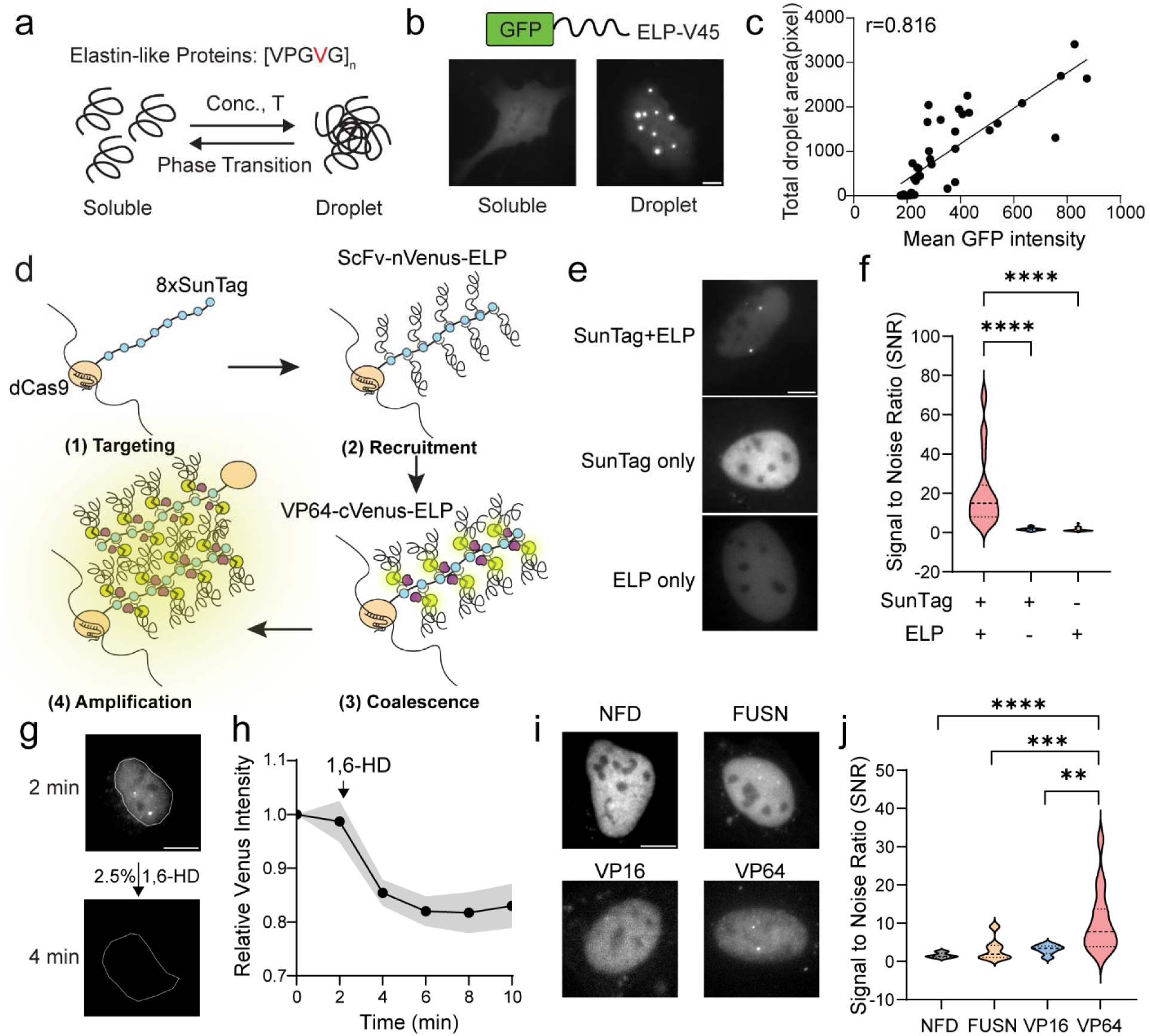
CRISPR-ELP enables robust, high-contrast condensate formation in live cells. **a.** Schematic of the elastin-like polypeptide (ELP) used in this study, composed of [VPGVP] pentapeptide repeats. ELPs undergo reversible transitions between a soluble random-coil state and a droplet state depending on temperature (T) and concentration (Conc.). **b.** ELP-V45 is tagged with GFP at the N-terminal and transfected into HEK 293T cells. Both phases of ELP can be observed in the cells depending on the ELP concentration. Scale bar, 10μm. **c.** Quantification of total droplet area as a function of GFP intensity for GFP–ELP-V45– expressing cells. r, Pearson correlation coefficient. n=39 cells. **d.** Schematic of the CRISPR-ELP droplet system. dCas9–8×SunTag is targeted to a genomic locus by a specific sgRNA (1. Targeting). ScFv–nVenus–ELP binds the SunTag repeats, locally enriching ELP (2. Recruitment), which triggers ELP-driven phase separation (3. Coalescence). Co-assembly of the split Venus halves and ELP droplets produces a bright, amplified fluorescent signal at the locus (4. Amplification). **e.** Representative images of SunTag-ELP, SunTag only or ELP only; in the SunTag only group, ScFv-eGFP was used; in the ELP only group, dCas9-ELP was used. Scale bar, 10μm. **f.** Quantification of signal to noise ratios (SNR) for the three groups shown in (e). n=31, 30, and 28 cells, respectively. Ordinary one-way ANOVA; **** p<0.0001. **g.** Representative images showing rapid dispersion of CRISPR-ELP droplets in U2-OS-SJ cells following treatment with 2.5% 1,6-hexanediol (1,6-HD). Dashed outlines indicate nuclei. Scale bar, 10μm. **h.** Quantification of the droplet intensity after 2.5% 1,6-HD treatment. n=5 cells. **i.** Representative images showing droplet formation for CRISPR-ELP constructs containing different functional domains. NFD, no-functional-domain group. Scale bar, 10 μm. **j.** Quantification of signal to noise ratios (SNR) for the functional domain variants shown in (i). n=16, 20, 6, and 15 cells, respectively. Ordinary one-way ANOVA; **** p<0.0001, *** p=0.0001, ** p=0.0086.

To enable robust visualization of non-repetitive genomic loci in live cells, overcoming the signal limitations ^11–15^, we developed a multi-component CRISPR-guided ELP droplet system (**Figure 1d**). The system comprises: **(1)** a catalytically inactive Cas9 (dCas9) fused to eight SunTag peptide repeats (dCas9-8xSunTag), directed to the genomic target by a specific guide RNA (gRNA); **(2)** a recruiting protein, ScFv-nVenus-ELP, where the ScFv antibody fragment binds to the SunTag repeats, tethering the N-terminal half of the Venus fluorescent protein (nVenus) and an elastin-like polypeptide (ELP) variant to the dCas9 scaffold; **(3)** a second ELP-containing protein, VP64-cVenus-ELP, which includes a VP64 transcriptional activation domain, the C-terminal half of Venus (cVenus), and a compatible ELP variant. The dCas9-SunTag platform acts as a nucleation site by recruiting multiple ScFv-nVenus-ELP molecules, thereby creating a high local copy number of ELP. The co-expressed VP64-cVenus-ELP is then expected to co-accumulate at this site, driven by ELP-ELP interactions to recruit more copies for the multiplex assembly and co-phase separation. The use of split Venus (nVenus and cVenus) is intended to ensure that fluorescence primarily arises upon the co-assembly of both ELP-containing components, thereby minimizing background signals from unbound precursors ^40^. The VP64 domain was included for its known capacity to interact with transcriptional machinery, potentially stabilizing the condensate and enabling transcriptional manipulation at the target site ^41,42^; **(4)** the combined LLPS of both ELP-containing proteins, and the stabilization of the functional domain VP64, is designed to generate a highly amplified fluorescent signal, forming a visible droplet at the target locus with a potential to regulate genome activities.

With all components co-expressed, the complete CRISPR-ELP system successfully generated bright nuclear droplets (**SunTag+ELP, Figure 1e,f**). To dissect which elements were necessary for this behavior, we removed each element from the full system to examine the effect. A SunTag-only reporter in which ELP modules were replaced with ScFv-GFP produced no significant puncta (**SunTag only, Figure 1e,f**), indicating that SunTag-mediated recruitment alone is insufficient to nucleate condensates. Conversely, when the SunTag scaffold was omitted and only dCas9-ELP or EGFP-ELP was expressed, the formation of distinct nuclear droplets was impaired, demonstrating that ELP alone is insufficient and that SunTag-driven multivalent recruitment is required to locally concentrate ELP above the phase-transition threshold (**ELP only, Figure 1e,f**). Together, these controls show that neither recruitment alone nor ELP phase behavior alone is adequate; robust coalescence requires the synergistic combination of SunTag-mediated clustering and ELP-driven LLPS. Consistent with their LLPS origin, the CRISPR-ELP droplets were also sensitive to 1,6-hexanediol treatment, which can disperse the condensates (**Figure 1g,h**).

To further investigate the contribution of the functional domain within the VP64-cVenus-ELP component, we replaced VP64 with other domains or removed it entirely. We tested: the potent transcriptional activator VP16; the N-terminal low complexity domain of the Fused in Sarcoma protein FUSN, known for its role in LLPS and ribonucleoprotein granule formation ^43,44^; and a ‘no-functional-domain’ version. A clear dependence on the nature of the functional domain was observed. Robust droplet formation was primarily supported only when VP64 was part of the cVenus-ELP construct. In contrast, when FUSN, VP16 or ‘no functional domain’ was present in the cVenus-ELP molecule, the formation of droplets was significantly impaired under the same conditions (**Figure 1i, j**). These findings indicate that the functional domain of VP64 on the second ELP protein plays a critical role, likely by stabilizing or enhancing the condensate through interactions with local cellular machinery.

### 2. CRISPR-ELP droplets specifically label non-repetitive genomic loci

To validate the specificity of CRISPR-ELP droplets, we employed the U2-OS-SJ cell line, which carries multiple insertions of synthetic TetO repeats in the chromosome 1p36 region and can be robustly labeled by TetR-HaloTag ^45,46^. This system provides a well-defined chromosomal landmark that can be used as a reference to evaluate targeting precision. Around the same 1p36 region, we chose a non-repetitive DNA sequence as the target site for the CRISPR-ELP droplet system (**Figure 2a**). When the CRISPR-ELP system was expressed in cells together with the targeting sgRNA, we consistently observed two discrete nuclear puncta, corresponding to labeling of the two alleles present on homologous chromosomes (**Figure 2b, arrows**). Because the engineered TetO repeats are integrated into only one of the two homologous chromosomes in the U2-OS-SJ cell line, only one of the two CRISPR-ELP droplets co-localized with the TetR-HaloTag (JF646) reference signal. This co-localization pattern is consistent with the known genomic configuration and supports that CRISPR-ELP droplets specifically label the intended non-repetitive locus. In contrast, cells lacking the targeting sgRNA showed no consistent co-localization (**Figure 2b**). To further validate the spatial relationship between the CRISPR-ELP droplet and the targeted genomic locus with higher precision, we performed DeepSIM super-resolution imaging followed by three-dimensional reconstruction. A line was first drawn across the droplet, and fluorescence intensity profiles were extracted along this line. The CRISPR-ELP and TetR-HaloTag signals exhibited correlated intensity peaks, indicating precise spatial alignment (**Figure 2c**). Zoomed-in views of the reconstructed volume, shown as orthogonal xy and xz projections, revealed tight spatial overlap between the CRISPR-ELP signal and the reference locus in all dimensions, confirming precise colocalization at the targeted genomic site (**Figure 2d**). These data further demonstrate that the CRISPR-ELP droplet forms in direct proximity to the chromatin locus rather than merely appearing colocalized in two-dimensional widefield images. We quantified locus-specific droplet assembly by measuring the enrichment of CRISPR-ELP droplet fluorescence at the reference site, using a circular ROI centered on the reference punctum and normalizing ELP intensity to a local background annulus. Targeting gRNAs produced strong ELP enrichment at the reference locus, indicating specific labeling of CRISPR-ELP droplets (**Figure 2e**).

**Figure 2.**
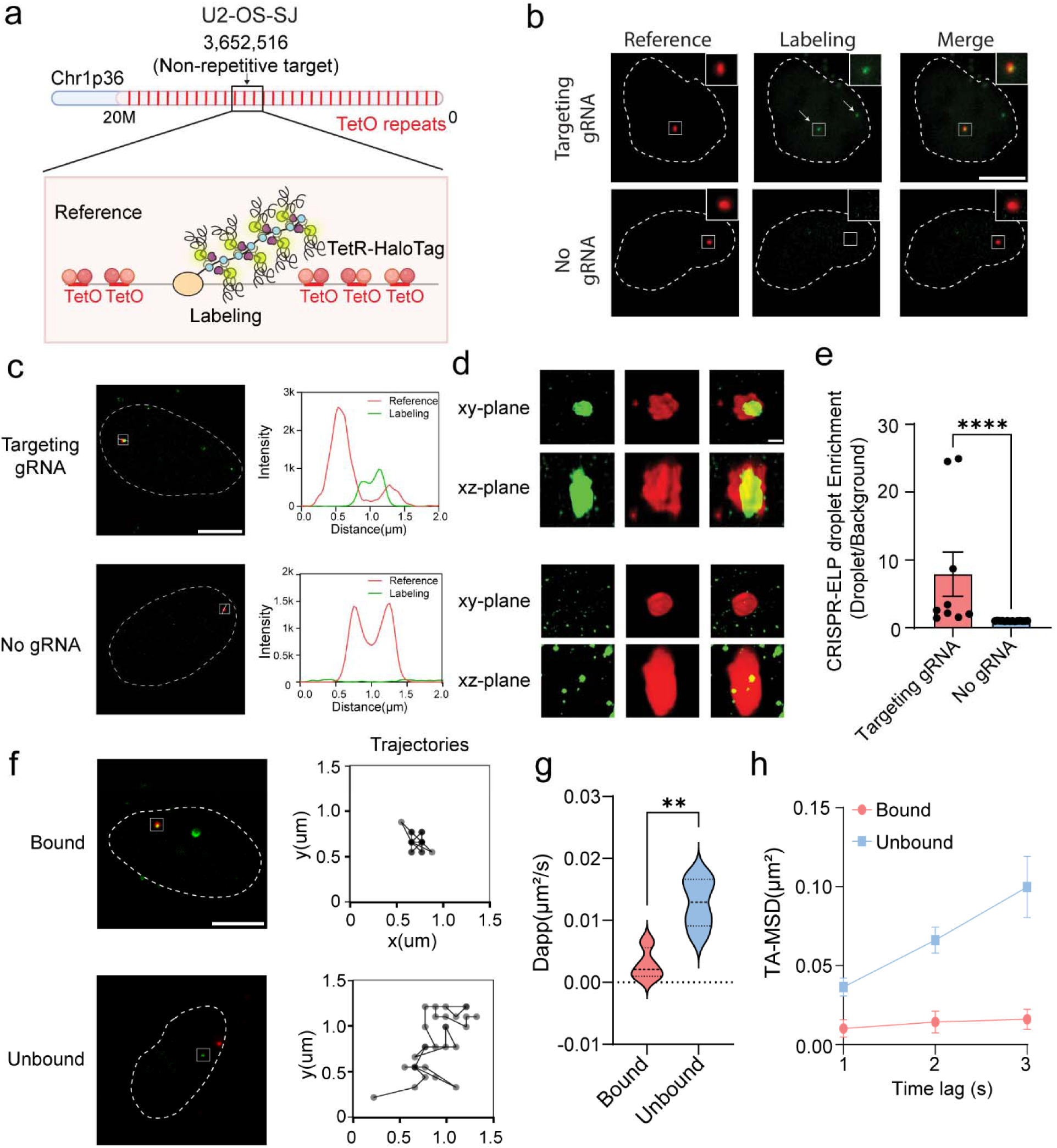
CRISPR-ELP droplets specifically label non-repetitive genomic loci. **a.** Schematic of the genomic targeting strategy in U2-OS-SJ cells. Multiple TetO repeats are integrated at chromosome 1p36 and serve as a reference labeled by TetR-HaloTag, while a nearby non-repetitive sequence within the same region is targeted by the CRISPR-ELP system. **b.** Representative widefield fluorescence images of U2-OS-SJ cells showing the TetR-HaloTag reference signal (red), CRISPR-ELP labeling signal (green), and merged channels. Top row, cells expressing the targeting sgRNA; bottom row, cells without sgRNA. Dashed lines outline the nuclear boundary. White arrows indicate CRISPR-ELP puncta on homologous alleles. Boxed regions indicate areas shown at higher magnification in the insets. Scale bar, 10 μm. **c.** DeepSIM super-resolution image of the CRISPR-ELP droplet and TetR-HaloTag reference. A line drawn across the droplet–locus structure (dashed line) was used to extract fluorescence intensity profiles shown on the right. Scale bar, 10 μm. **d.** Orthogonal xy and xz projections of the three-dimensional DeepSIM reconstruction corresponding to the region analyzed in (c). Scale bar, 2μm. **e.** Quantification of locus-specific CRISPR-ELP enrichment. Fluorescence intensity of the CRISPR-ELP signal was measured within a circular ROI centered on the TetR reference punctum and normalized to a surrounding background annulus. N=9 and 14 cells for the targeting gRNA and no gRNA groups respectively. Mann-Whitney test; **** p<0.0001. **f.** Representative trajectories of CRISPR-ELP puncta tracked over 15 seconds in the presence (bound) or absence (unbound) of the targeting sgRNA, illustrating confined versus diffusive motion, respectively. Scale bar, 10 μm. **g.** Apparent diffusion coefficients estimated from the shortest time lag of mean-squared displacement (MSD) analysis. n=6 and 4 cells for bound and unbound groups. Mann-Whitney test; ** p=0.0095. **h.** Time-averaged MSD (TA-MSD) curves for bound and unbound CRISPR-ELP puncta. n=6 and 4 cells for bound and unbound groups.

Because both systems (TetR-HaloTag and CRISPR-ELP droplet) are present in the same cell, we next evaluated whether the CRISPR-ELP droplet might be indirectly recruited by TetR bound to TetO, rather than directly recognizing the DNA sequence defined by the sgRNA. To exclude this possibility, cells were pre-cultured in medium containing 500 nM doxycycline to block TetR binding to TetO sites. Under these conditions, TetR-HaloTag became diffuse and no longer marked TetO arrays, whereas CRISPR-ELP droplets still formed at specific nuclear locations (**Supplementary Figure S3a**). Following doxycycline washout, TetR-HaloTag reappeared at the TetO repeats locations and regained co-localization with pre-existing CRISPR-ELP droplets. This temporal sequence demonstrates that CRISPR-ELP droplets localize independently of TetR binding, and co-localization reflects their convergence on the same chromosomal region rather than artificial recruitment.

In the absence of gRNA, we occasionally observed small, dim CRISPR-ELP puncta. We analyzed their dynamic behavior using time-lapse imaging and particle tracking and compared droplet mobility with or without gRNA. On-target CRISPR-ELP droplets remained confined to a stable nuclear position over the imaging period, consistent with being tethered to chromatin at a fixed locus. In contrast, puncta observed in the no-sgRNA group were highly mobile, exhibited rapid diffusive motion (**Figure 2f**). Mean-squared displacement (MSD) analysis revealed markedly reduced mobility of bound droplets compared to unbound droplets. The apparent diffusion coefficient estimated from the shortest time lag was approximately 4.5-fold higher for unbound droplets than for bound droplets (0.013 μm²/s versus 0.0029 μm²/s) (**Figure 2g**). Consistent with this difference, time-averaged MSD (TA-MSD) curves of bound droplets increased more slowly with time lag than those of unbound droplets, indicating constrained motion upon binding (**Figure 2h**). These differences in mobility and diffusion behavior provide a robust set of metrics for further distinguishing specific versus nonspecific CRISPR-ELP puncta. Together, these results show that CRISPR-ELP droplets can robustly and specifically label endogenous non-repetitive loci in living cells with sufficient brightness and positional stability for downstream functional analysis. This capability provides a foundation for coupling locus visualization to dynamic measurements of DNA repair at defined genomic sites. Together, these results establish that CRISPR-ELP droplets can specifically label endogenous non-repetitive loci in live cells, providing the basis for integrating locus visualization with functional measurements of DNA repair.

### 3. GLIDRE integrates locus labeling, DSB induction and live-cell monitoring of repair-associated dynamics

AsCas12a (Cpf1) is a commonly used CRISPR nuclease capable of inducing double-strand DNA breaks (DSBs) at gRNA-targeted sites. Unlike Cas9, AsCas12a uses a completely different format of gRNA for targeting and mediating cutting, which makes it orthogonal and compatible with the dCas9-ELP mediated DNA labeling system ^47^. To monitor DSB repair at non-repetitive loci and simultaneously infer associated epigenetic modifications, we tested two well-characterized markers. The first, 53BP1, is a crucial DNA damage response protein whose recruitment and large-scale accumulation around DSB sites are dependent on chromatin ubiquitylation events, particularly H2AK15ub, thus serving as a dynamic indicator of this epigenetic signaling pathway ^48,49^. The second marker is the tandem BRCT domain from MDC1 (Mediator of DNA Damage Checkpoint 1), which directly binds to phosphorylated histone H2AX (γH2AX), a primary epigenetic mark rapidly established at DSB sites ^50^. For proof of concept, we first targeted a non-repetitive site within the chr1p36 region (pre-labeled with TetR-eGFP) for DSB induction by AsCas12a in U2OS-SJ cells (**Supplementary Figure S4a**). The subsequent recruitment of ectopically expressed 53BP1-HaloTag or MDC1 BRCT-HaloTag was observed under microscope. Both markers successfully accumulated at the induced DSB sites in live cells, indicating their utility in visualizing repair foci and their associated epigenetic contexts (**Supplementary Figure S4b**). 53BP1-HaloTag was selected for its clearer signal and established utility in robustly detecting DSB sites for subsequent GLIDRE studies, where its dynamics would reflect the progression of ubiquitination-dependent repair pathways (**Supplementary Figure S4c**).

To precisely control the timing of DSB, we generated a stable U2OS-SJ cell line that contains a heat-inducible AsCas12a gene cassette under the synthetic promoter 7HE-YB (U2OS-SJ-iCas12a) ^51^ so that we can control the starting time of DNA repair (**Supplementary Figure S5a**). Upon heat shock at 43°C for 20 minutes, AsCas12a expression was efficiently induced, reaching peak levels six hours post-heat treatment (**Supplementary Figure S5b**). The expressed AsCas12a, guided by its specific gRNA, is expected to introduce targeted DSBs at the designated DNA site, enabling precise temporal control over DSB induction. We then targeted a locus within the 1p36 region using TetR-GFP and tested the DSB repair induced by the heat-inducible AsCas12a. We are able to capture the process of DSB repair in live cells, and 53BP1 nicely tracked and inferred the dynamics of DSB repair process (**Supplementary Figure S5c and Supplementary video 1**).

To achieve simultaneous visualization of non-repetitive loci and DSB repair dynamics, including its associated epigenetic context, we developed the GLIDRE (<u>G</u>enomic <u>L</u>ocus Imaging and <u>D</u>SB <u>R</u>epair Monitoring by <u>E</u>lastin-like Proteins) system (**Figure 3a**). This integrated design extends locus imaging beyond positional tracking by enabling simultaneous locus labeling, controlled DSB induction, and real-time monitoring of repair-associated chromatin dynamics at the same genomic site. In this approach, CRISPR-ELP droplets dynamically label the target DNA region. Concurrently, DSBs are induced adjacent to this labeled site using heat-induced AsCas12a, and the subsequent repair process is monitored using 53BP1-HaloTag, whose recruitment reflects ongoing chromatin ubiquitylation and machinery assembly events essential for this repair pathway. This configuration offers precise spatiotemporal control over DNA damage induction and allows concurrent, real-time monitoring of both the specific genomic locus and the associated epigenetic events during repair response.

**Figure 3.**
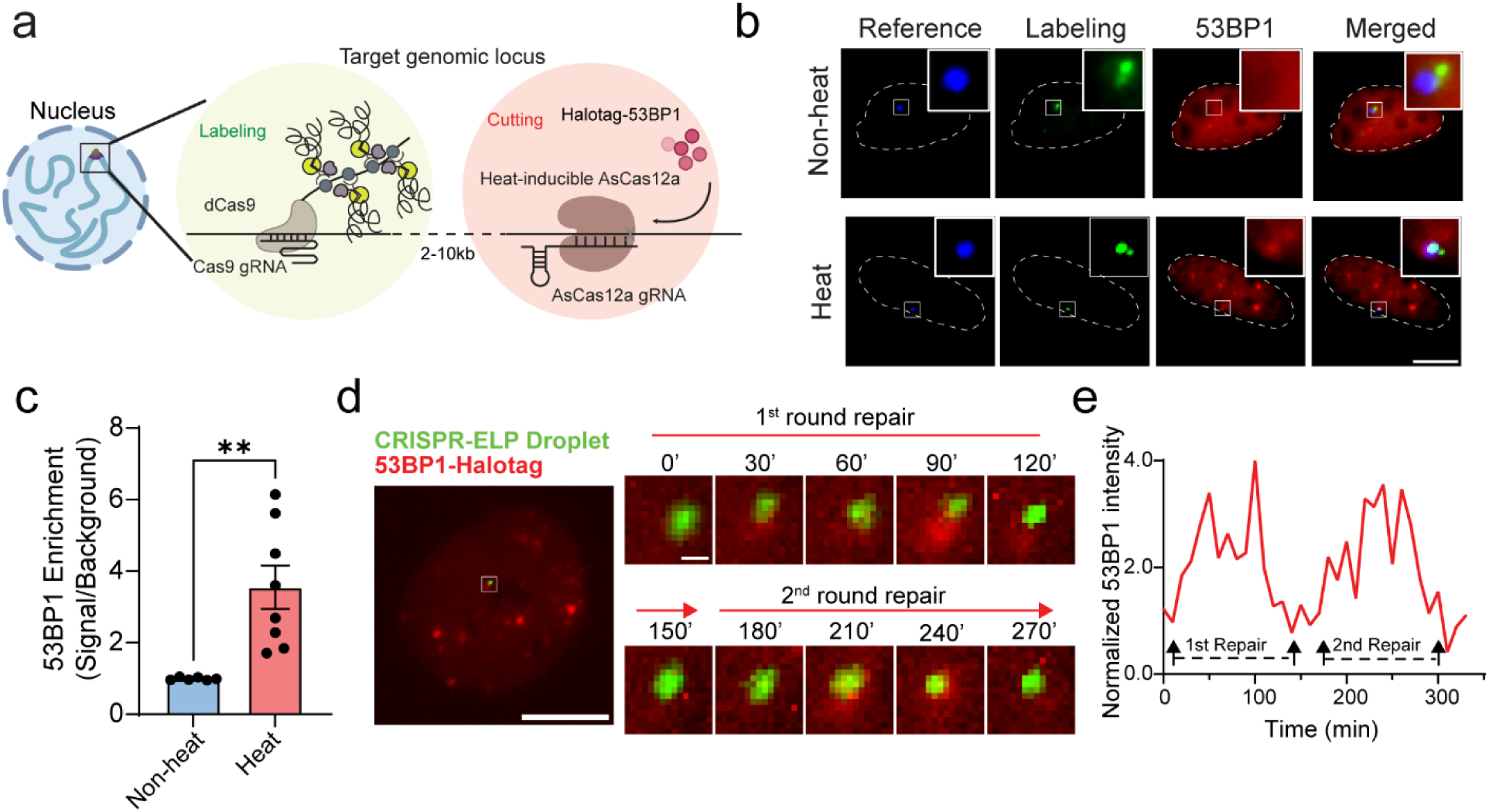
Genomic Locus Imaging and DSB Repair Monitoring by Elastin-like Proteins (GLIDRE) **a.** Schematic of the GLIDRE platform for concurrent genomic locus labeling and double-strand break (DSB) monitoring. In the zoomed-in view, the target locus is labeled by a CRISPR-ELP droplet (left), while a nearby DSB site is induced by orthogonal, heat-inducible AsCas12a and monitored by HaloTag-labeled 53BP1 (right). The labeled locus and the AsCas12a cutting site are positioned approximately 2–10 kb apart. **b.** Representative live-cell images showing GLIDRE imaging at a non-repetitive target locus in the chr1p36 region under non-heat and heat conditions. Columns show the chromosomal reference (TetR-BFP, blue), CRISPR-ELP droplet signal (green), 53BP1-HaloTag signal (red), and merged images. Insets show enlarged views of the boxed regions. Scale bar, 10 μm. **c.** Quantification of 53BP1 enrichment (signal/background) at the GLIDRE-labeled locus under non-heat and heat conditions. n=6 and 8 for non-heat and heat groups respectively. Welch’s t-test; ** p=0.0089. **d.** Representative time-lapse images showing repair-associated 53BP1 dynamics at a CRISPR-ELP droplet-labeled locus on chromosome 2. Left, whole-cell view with the tracked locus indicated by a box. Right, enlarged views of the target region over time. Green, CRISPR-ELP droplet; red, 53BP1-HaloTag. Two episodes of 53BP1 recruitment and dissociation are observed (annotated as the 1st and 2nd round repair). Scale bar, 10 μm. **e.** Time course of normalized 53BP1 signal intensity at the CRISPR-ELP droplet-labeled locus corresponding to the cell shown in (d).

We first validated the GLIDRE system by targeting a non-repetitive locus in the chr1p36 region. Here, TetR-tBFP (blue) served as the chromosomal reference, CRISPR-ELP droplets (green) provided dynamic visualization of the locus, and 53BP1 (red) tracked the DSB repair foci and its associated epigenetic signals. Without heating, we observed colocalization between reference TetR-BFP and CRISPR-ELP droplet, but no 53BP1 signal was detected to co-localize with them. After heating, clear colocalization of TetR, CRISPR-ELP droplets, and the 53BP1 repair/ubiquitination signal was observed, suggesting the heat inducibility of the system (**Figure 3b**). Quantitative analysis revealed that cells exhibiting colocalization between the CRISPR-ELP droplets and the 53BP1 signal also showed increased 53BP1 signals aligning with the TetR reference channel, highlighting the high precision of GLIDRE in integrating locus visualization with the monitoring of site-specific repair and epigenetic events (**Figure 3c**).

We further utilized the GLIDRE system to dynamically track the repair process and its associated epigenetic dynamics at another non-repetitive locus outside the chr1p36 region. Using time-lapse imaging, we captured multiple rounds of recruitment and dismissal of 53BP1 at the target site, reflecting possibly multiple rounds of DSB repairs (**Figure 3d and supplementary video 2**). The dwelling time of 53BP1-HaloTag at the DSB site provided a quantitative measure of the associated epigenetic kinetics of repair response at a specific genomic locus (**Figure 3e**). Together, these results underscore the utility of GLIDRE in enabling precise, real-time analysis of locus specific epigenetic dynamics associated with DNA repair within living cells.

### 4. GLIDRE identifies chromatin accessibility as a dominant determinant of DNA double-strand break repair dynamics

Recent studies have highlighted transcriptional activity as a potential modulator of DNA repair efficiency ^52^. In fact, evidence from fungal models has demonstrated that actively transcribed genes repair DSBs more efficiently than their inactive counterparts ^53^. We directly investigated the impact of local transcriptional activity on DSB repair in live mammalian cells, utilizing our GLIDRE system. A key design feature of GLIDRE is the incorporation of the potent transcriptional activator VP64 within the CRISPR-ELP droplet, enabling us to precisely induce and control transcription at targeted genomic loci while concurrently monitoring DSB repair dynamics. By varying the proximity of CRISPR-ELP droplets to the transcription start site (TSS) of the targeted genes, we could alter the transcription levels of the target genes. As an initial model, we selected IL-1B, a pro-inflammatory gene known to undergo dynamic chromatin remodeling and inducible accessibility upon stimulation ^54,55^. Positioning the CRISPR-ELP droplets close to the TSS of the IL-1B gene resulted in robust transcriptional upregulation, elevating expression levels by several thousand-fold (**Figure 4a,b**). Comparing 53BP1 dwelling time at these different transcription levels revealed a significant enhancement in repair efficiency when the IL-1B gene was actively transcribed. Specifically, the 53BP1 dwell time for DSBs was nearly three times as faster in the transcriptionally active state (∼110min) compared to the inactive state (∼340min) (**Figure 4c-e**). This finding provides the first direct evidence, in live mammalian cells, of transcription-coupled DSB repair, supporting the hypothesis that transcriptional activity can affect DSB repair efficiency.

**Figure 4.**
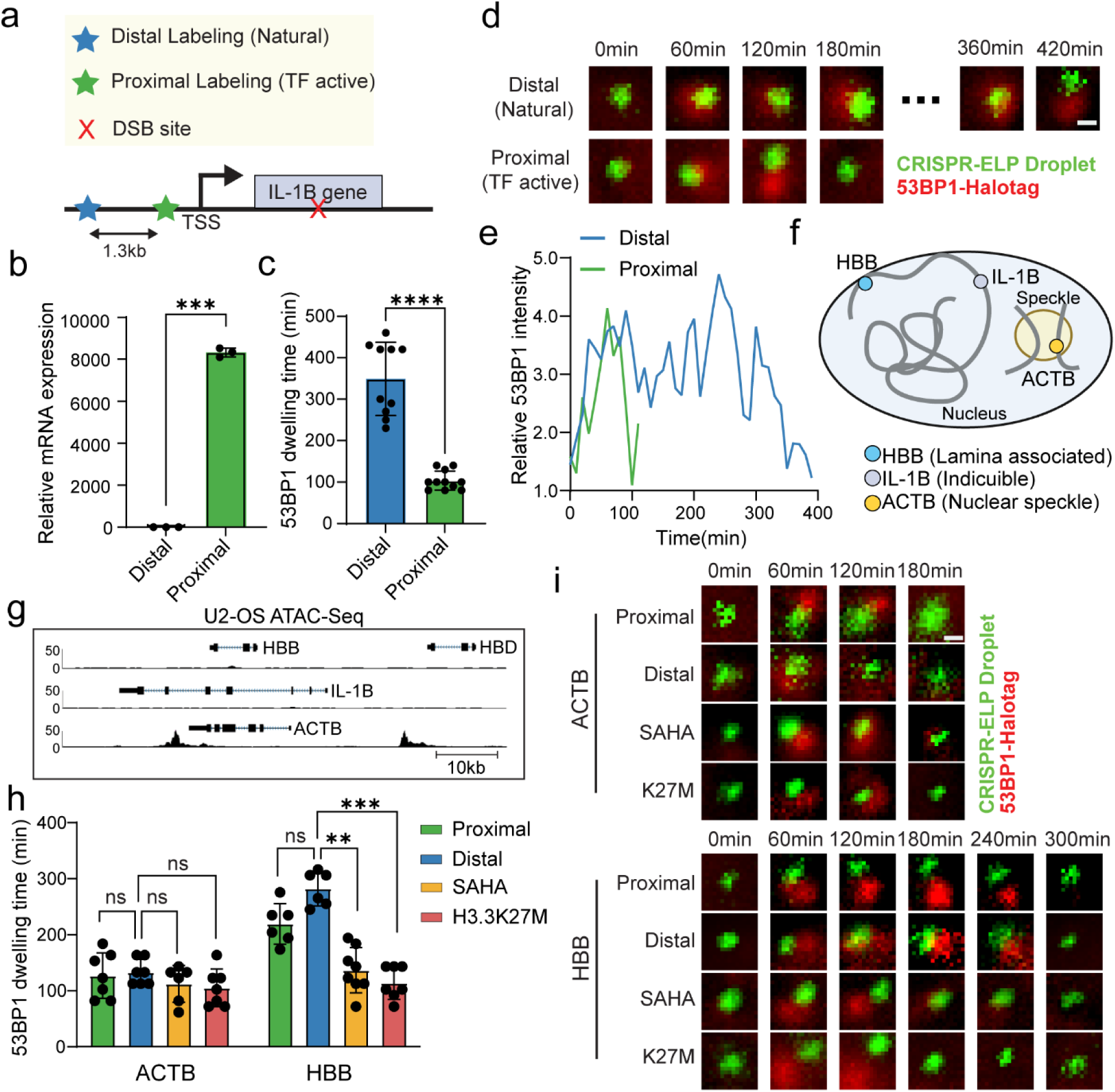
Chromatin accessibility defines the baseline for locus-specific DSB repair-associated dynamics. **a.** Experimental design for transcription modulation at the IL-1B locus using GLIDRE. CRISPR-ELP droplets were positioned either distal to the transcription start site (TSS) (“Distal labeling” natural transcription) or proximal to the TSS (“Proximal labeling” VP64 transcription factor active), while the AsCas12a DSB site (red X) was placed within the gene body. **b.** RT–qPCR quantification of IL1B mRNA under distal versus proximal labeling conditions. Bars show mean ± SD; n=3 independent biological repeats. Welch’s t-test; *** p=0.0002. **c.** Quantification of 53BP1 dwell time (repair-associated signaling duration) at the IL-1B DSB site between distal versus proximal labeling conditions. Bars show mean ± SD; n=10 and 11 nuclei from at least 3 independent biological repeats for distal and proximal labeling, respectively. Mann-Whitney test; **** p<0.0001. **d.** Representative time course images of GLIDRE at the IL1B locus comparing distal (natural) and proximal (TF active) conditions. Green, CRISPR-ELP droplet; red, 53BP1-HaloTag. Scale bar, 0.5μm. **e.** Representative time course of relative 53BP1 intensity at the GLIDRE-labeled IL1B locus for distal (blue) and proximal (green) conditions, corresponding to the example in (d). **f.** Schematic illustrating the three loci analyzed for chromatin context: HBB (lamina-associated/compact), IL1B (inducible), and ACTB (housekeeping, nuclear speckle associated), with their typical nuclear environments. **g.** Genome browser tracks showing U2-OS ATAC-seq peaks at the HBB, IL1B, and ACTB loci (GSE139190). Scale bar, 10 kb. **h.** Summary quantification of 53BP1 dwell time at ACTB and HBB under the indicated conditions: Proximal (TF active), Distal (natural), SAHA treatment, and H3.3K27M expression. Bars show mean ± SD; n=6-7 nuclei from at least 3 independent biological repeats for each groups. Statistical analysis are conducted separately for ACTB and HBB groups. Kruskal-Wallis test; ** p=0.0049, *** p=0.0004. **i.** Representative time-lapse images showing GLIDRE dynamics at ACTB (top) and HBB (bottom) under the indicated conditions (Proximal, Distal, SAHA, and H3.3K27M). Green, CRISPR-ELP droplet; red, 53BP1-HaloTag. Scale bar, 0.5μm.

Given that transcription may remodel chromatin structure by shifting regions of tightly packed heterochromatin toward more accessible euchromatin, we examined whether transcription is a key determinant of DSB repair efficiency independent of chromatin architecture ^56^. Chromatin exists in distinct states, with heterochromatin characterized by densely packed, transcriptionally repressive regions, and euchromatin by more accessible, transcriptionally active regions ^57^. To examine the interplay between chromatin status and DSB repair, we selected two additional genes for analysis: HBB, associated with the nuclear lamina and representative of a tightly regulated locus often found in a compacted, heterochromatic state in non-erythroid cells ^58,59^, and ACTB, a ubiquitously expressed housekeeping gene and strongly associated with nuclear speckle, generally residing in open, euchromatic regions ^60^ (**Figure 4f,g and Supplementary Figures S6a,b**).

In initial experiments, we measured 53BP1 dwell time under natural transcription levels by positioning CRISPR-ELP droplets far away from the TSS of each gene (**Supplementary Figure S7a**). The results revealed that repair at ACTB, located in an open chromatin region, was significantly faster (∼130min), nearly three-fold compared to both IL-1B (∼350min) and HBB (∼280min) (**Figure 4h,i**). To further dissect the interplay between induced transcription and basal chromatin state, we again leveraged the integrated VP64 activator in GLIDRE. By repositioning CRISPR-ELP droplets closer to the TSS of ACTB and HBB, we induced transcriptional activation at both loci (**Supplementary Figure S7b**). Although ACTB transcription increased, DSB repair dynamics were not significantly altered. ACTB is already actively transcribed and proximal targeting produced only a modest (∼1.5-fold) increase in expression (**Figure 4h,i and Supplementary Figure S7b**). Notably, despite robust transcriptional activation at HBB(∼100-fold), DSB repair dynamics also remained largely unchanged (**Figure 4h,i**). Together, these findings suggest that basal chromatin accessibility may dominate DSB repair dynamics, whereas transcriptional activation influences repair most strongly at loci with more plastic, inducible chromatin states, such as IL-1B ^55^.

To test the role of chromatin accessibility more directly, we treated cells with SAHA, a histone deacetylase (HDAC) inhibitor known to increase histone acetylation and promote chromatin relaxation ^61^. Following SAHA treatment, HBB exhibited accelerated DSB repair kinetics, despite minimal changes in transcriptional activity (**Figure 4h-i and Supplementary Figure S7c**). To reduce the possibility that SAHA accelerated DSB repair through mechanisms unrelated to chromatin opening, we additionally overexpressed the oncohistone H3.3K27M, which is associated with increased chromatin accessibility (**Supplementary Figure S8**). Consistent with the SAHA results, H3.3K27M expression also accelerated DSB repair kinetics at HBB with minimal changes in transcriptional output (**Figure 4h,i and Supplementary Figure S7c)**. Together, these data support a hierarchical model in which chromatin accessibility defines the baseline capacity for DSB repair dynamics, whereas transcriptional activity modulates these dynamics primarily within permissive or plastic chromatin contexts.

### 5. H3.3K27M reshapes locus-specific DNA repair dynamics associated with tumor resistance

We next asked whether locus-specific repair-associated dynamics might contribute to tumor resistance in a chromatin-deregulated setting. Because chromatin accessibility emerged as a major determinant of repair-associated dynamics in the preceding analyses, we reasoned that oncogenic chromatin remodeling could reshape genome maintenance states at selected loci and thereby influence tumor survival under therapeutic stress. H3.3K27M provides a compelling model for this question, as it drives broad chromatin reprogramming and is strongly associated with treatment resistance in diffuse intrinsic pontine glioma (DIPG) ^62–64^. To model these effects in a tractable system, we used U2-OS cells, which have been employed previously to study H3.3K27M-driven chromatin phenotypes and to recapitulate key features reported in DIPG models ^65,66^. Consistent with the therapeutic resistance phenotype, H3.3K27M-expressing U2-OS cells exhibited increased survival compared to H3.3K27WT-expressing control cells when subjected to genotoxic stress induced by telomere shortening via the CRISPR-Cas9 system ^67,68^ (**Supplementary Figure S9a,b**).

CAR T-cell cytotoxicity can impose severe stress on target tumor cells and has been reported to engage DNA damage responses ^69,70^. To model CAR T responses in our system, we generated PD-L1-targeting CAR T-cells using high-affinity PD-1 as the recognition domain ^71^ (HACVPD-1; **Figure 5a**). CAR surface expression was confirmed by flow cytometry, and HACVPD-1 CAR T-cells exhibited strong cytotoxicity against PD-L1 positive U2-OS cells in 2D co-culture (**Supplementary Figure S10a,b**). Using this validated CAR T platform, we found that spheroids generated from H3.3K27M-expressing cells were significantly more resistant to HACVPD-1 CAR T-mediated killing than H3.3K27WT spheroids (**Figure 5b,c**). H3.3K27M is known to reduce global H3K27me3 levels while redistributing Polycomb activity to specific genomic loci ^72^. We therefore hypothesized that H3.3K27M-driven redistribution of the chromatin landscape creates a locus-specific imbalance in genome maintenance, in which some regions become more permissive to repair (such as IL1B and HBB in Figure 4), whereas other regions, particularly key tumor suppressor loci, become more vulnerable to genotoxic stress. Examination of previously published H3K27me3 ChIP-seq and ATAC-seq datasets showed that the tumor suppressor gene CDKN2A exhibits markedly increased H3K27me3 together with limited chromatin accessibility in H3.3K27M cells relative to control cells (**Figure 5d**) ^73^.

**Figure 5.**
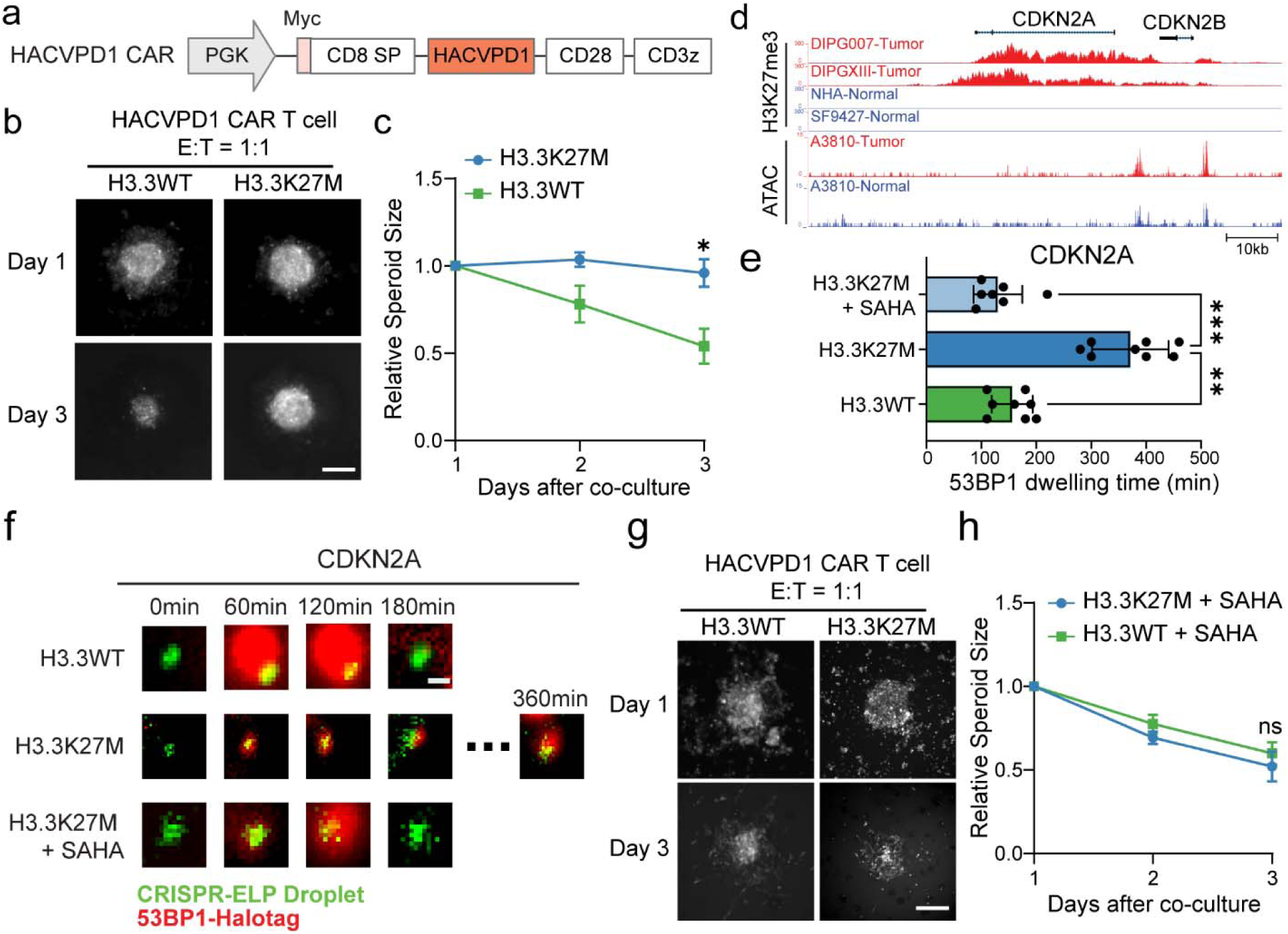
H3.3K27M reshapes locus-specific DSB repair dynamics in association with resistance and SAHA response. **a.** Schematic of the HACVPD-1 CAR construct used in this study. The CAR is expressed under the PGK promoter and contains a Myc tag, CD8 signal peptide (CD8 SP), HACVPD-1 binding domain, CD28 costimulatory domain, and CD3ζ signaling domain. **b.** Representative images of tumor spheroids formed from H3.3WT or H3.3K27M cells during co-culture with HACVPD-1 CAR T-cells (E:T = 1:1) at Day 1 to Day 3. Scale bar, 200 μm. **c.** Quantification of relative spheroid size over time following co-culture with HACVPD-1 CAR T-cells (E:T = 1:1). Bars show mean ± SEM; n= 4 spheroids. Statistical analysis is performed for Day 3. Unpaired student t-test; * p=0.0169. **d.** Genome browser tracks showing H3K27me3 and ATAC-seq signals at the CDKN2A region across DIPG and control datasets. Tracks shown for DIPG007, DIPGXIII, NHA, SF9427 (H3K27me3, GSE162976) and A3810-Tumor vs A3810-Normal (ATAC, GSE162976). Tracks are generated using UCSC genome browser (http://genome.ucsc.edu). Scale bar, 10 kb. **e.** GLIDRE quantification of DSB repair-associated dynamics at the CDKN2A locus in H3.3WT, H3.3K27M, and H3.3K27M + SAHA conditions. Bars show mean ± SD; n=7-8 nuclei from at least 3 independent biological repeats. Kruskal-Wallis test; *** p=0.0005, ** p=0.0089. **f.** Representative time course images of GLIDRE at the CDKN2A locus comparing H3.3WT, H3.3K27M, and H3.3K27M + SAHA conditions. Green, CRISPR-ELP droplet; red, 53BP1-HaloTag. Scale bar, 0.5μm. **g.** Representative images of tumor spheroids after SAHA pretreatment followed by co-culture with HACVPD-1 CAR T-cells (E:T = 1:1) at Day 1 to Day 3 for H3.3WT and H3.3K27M spheroids. Scale bar, 200 μm. **h.** Quantification of relative spheroid size over time following treatment with SAHA and co-culture with HACVPD-1 CAR T-cells (E:T = 1:1). Bars show mean ± SEM; n= 6 spheroids. Statistical analysis is performed for Day 3. Unpaired student t-test; ns p=0.4925.

We therefore focused on CDKN2A as a representative repressed tumor suppressor locus to test whether H3.3K27M-associated chromatin changes alter locus-specific DSB repair dynamics. We first used GLIDRE to measure DSB repair-associated dynamics at the CDKN2A locus in H3.3K27WT-and H3.3K27M-expressing cells. To minimize perturbation of basal transcription, GLIDRE labeling was positioned distal to CDKN2A’s TSS. In H3.3K27M**-**expressing cells, 53BP1 dwelling time (∼370min) was significantly prolonged relative to H3.3K27WT cells (∼160min), indicating delayed repair-associated dynamics at this locus (**Figure 5e,f**). The CDKN2A phenotype is also consistent with reports that PRC2 and H3K27 methyltransferase activity can be aberrantly enriched at specific loci, including tumor suppressor genes such as CDKN2A, in H3.3K27M backgrounds ^72^. Together, these findings support a model in which H3.3K27M redistributes repair capacity across the genome, promoting faster repair at some loci while impairing repair at tumor suppressor loci such as CDKN2A. Because CDKN2A is a critical regulator of tumor suppression, impaired maintenance of this locus under genotoxic stress may further weaken growth-restrictive programs and thereby contribute to the therapy-resistant state of H3.3K27M tumors.

We next asked whether modulating chromatin accessibility could alter locus-specific DSB repair and sensitize H3.3K27M cells to CAR T-cell therapy. We treated cells with SAHA, a histone deacetylase (HDAC) inhibitor that increases histone acetylation and promotes chromatin relaxation. Using GLIDRE, we measured DSB repair-associated dynamics at the CDKN2A locus in H3.3K27M cells with SAHA treatment and found that SAHA significantly accelerated repair at CDKN2A (∼130min) (**Figure 5e,f**). These data support the interpretation that SAHA relieves local chromatin repression and restores CDKN2A repair dynamics toward the WT state. Accordingly, we pretreated tumor spheroids (H3.3K27WT and H3.3K27M) with SAHA prior to co-culture with HACVPD-1 CAR T-cells. SAHA pretreatment sensitized H3.3K27M spheroids to CAR T therapy, with no statistically significant difference detected compared to H3.3WT spheroids (**Figure 5g,h**). Together, these results suggest that chromatin modulation by SAHA shifts H3.3K27M cells toward a more WT-like response to CAR T-cell therapy. CDKN2A provides a representative example of how H3.3K27M-associated chromatin changes alter repair-associated dynamics at specific gene loci and are associated with the resistant phenotype.

## Discussions

In this study, we introduced the GLIDRE system, a novel live-cell imaging platform that leverages CRISPR-guided ELP droplets to visualize DNA double-strand break repair at, in principle, any genomic loci in real time. A major advantage of GLIDRE is that it extends locus imaging beyond genomic labeling by integrating locus visualization, controlled DSB induction, and live-cell repair tracking in a single system. As such, our method offers unique opportunities to reveal the spatiotemporal landscape of genomic DNA repair, including the recruitment of key repair factors and the progression of associated epigenetic modifications.

Using GLIDRE, we found that chromatin accessibility at specific loci strongly influences DSB repair efficiency. Specifically, we observed that open chromatin undergoes faster repair, whereas compacted genomic loci exhibit slower repair kinetics, with transcriptional activity exerting more context-dependent effects. Our results support a hierarchical view of locus-specific DNA damage responses in which chromatin accessibility establishes a baseline repair-permissive state, while transcription modulates repair-associated dynamics chiefly at loci that are already permissive or plastic. The H3.3K27M cell model highlighted the functional significance of this regulation, as H3.3K27M caused a less accessible chromatin state at the tumor suppressor CDKN2A, thereby resulting in delayed DSB repair. These data suggest a mechanistic link between chromatin state and repair kinetics, consistent with prior studies showing that chromatin-associated factors can promote DNA repair by facilitating repair-factor access and recombination dynamics ^74^. Our findings suggest that tumor response to immunotherapy is shaped not only by classical immune evasion pathways, but also by chromatin-regulated genome maintenance states that influence how tumor cells tolerate therapy-induced stress. The therapeutic implications of this idea are underscored by our demonstration that chromatin modulation can alter DNA repair dynamics. Treatment with the HDAC inhibitor SAHA restored its repair dynamic, and sensitized H3.3K27M mutant cells to CAR T-cell therapy. These results indicate that SAHA treatment shifts H3.3K27M spheroids toward a WT-like response to CAR T-cell killing. This interpretation is also consistent with broader evidence that disruption or epigenetic silencing of tumor suppressor genes such as CDKN2A is associated with aggressive tumor behavior and poor therapeutic outcome, suggesting that impaired maintenance of such loci under genotoxic stress may further reinforce resistance ^75,76^. These results not only confirm that targeted epigenetic therapies can enhance the efficacy of immunotherapeutic strategies, but also provide a rationale for broader applications in oncology. For example, future studies could explore how modulation of chromatin accessibility might potentiate the effects of genotoxic agents in various cancer models, including those driven by other histone mutants or epigenetic deregulation.

In conclusion, the GLIDRE system advances the toolkit for studying genome dynamics by enabling precise, real-time visualization of DSB repair at endogenous, non-repetitive loci. Through its application, we have demonstrated the intricate relationship among transcription, chromatin accessibility, and DNA repair capacity, particularly in the context of oncogenesis and therapy resistance. Although the current implementation focuses on single-locus measurements using 53BP1 as a repair-associated readout, the platform should be extendable to additional chromatin and repair markers and to broader disease settings. By offering a flexible platform that can be adapted to study enhancer-promoter interactions, cell differentiation, and chromatin reorganization, this technology paves the way for a deeper understanding of genome biology and the design of novel therapeutic interventions targeting chromatin state and DNA repair pathways.

## Methods and materials

### Plasmids

Plasmids used in this study are listed in Supplementary Table 1. Plasmids were assembled using Gibson Assembly (NEB, E2611L) or T4 DNA ligation (NEB, M0202L). PCR amplification was performed using Q5 High-Fidelity DNA Polymerase (NEB, M0491). All constructs were verified by Sanger sequencing and Plasmid-EZ (Azenta Life Sciences).

ELP cloning: ELP variants (V10, V20, V30, V45) were PCR-amplified from pMAL-c5x-ELP[V60] (Addgene #6701) ^36^ using primers encoding appropriate restriction sites and assembled into expression backbones by T4 ligation.

gRNA sequences are used in this study are listed in Supplementary Table 2. gRNA sequence is designed using the online CRISPick tool from the Broad Institutue (https://portals.broadinstitute.org/gppx/crispick/public). For GLIDRE locus labeling, sgRNAs were selected to target non-repetitive regions proximal or distal to transcription start sites (TSS) as indicated in each figure. For DSB induction, AsCas12a crRNAs were designed to target a site 2–10 kb from the locus-labeling site (as indicated in Fig. 3–5).

HACVPD1 CAR construct is generated by replacing the the CD19 scFv of current 2^nd^ generation anti-CD19 CAR with the high-affinity PD1 extracellular domain ^77^.

### General cell culture

Human embryonic kidney cells (HEK293T) cells were cultured in Dulbecco’s modified Eagle’s medium (Gibco, 11995073) supplemented with 10% (vol/vol) heat-inactivated FBS (Gibco, 10438026), 100 units/mL penicillin-streptomycin (Gibco, 15140122). U2-OS-SJ cells were a gift from Dr. Xavier Darzacq at UC Berkely. These cells were cultured in McCoy’s 5A (Modified) Medium (Gibco, 16600082) supplemented with 10% (vol/vol) Tet system approved FBS (Takara, 631107), 100 units/mL penicillin-streptomycin. Human peripheral blood mononuclear cells (PBMCs) were isolated from buffy coats (San Diego Blood Bank, WB-PBCT-1BG-XXXX-02) by density gradient centrifugation using lymphocyte separation medium (Corning, 25-072-CV). Primary human T cells were then purified from the PBMC fraction using a Pan T Cell Isolation Kit (Miltenyi Biotec, 130-096-535) according to the manufacturer’s instructions. Purified T cells were cryopreserved in 92.5% fetal bovine serum (FBS) and 7.5% dimethyl sulfoxide (DMSO) until use. For culture, primary human T cells were maintained in X-VIVO 15 medium (Lonza, 04-418Q) supplemented with 5% FBS, 1% penicillin-streptomycin, 50 μM 2-mercaptoethanol (Gibco, 31350010), and 100 U/mL recombinant human IL-2 (PeproTech, 200-02). Cells were cultured in a humidified incubator at 37°C with 5% CO.

### Transfection

General transfection was performed using the Lipofectamine 3000 reagent (Invitrogen, L3000015) according to the manufacture’s protocol. In general, 0.3×10^6^ HEK 293T cells or 0.1×10^6^ U2-OS-SJ cells were seeded on 12-well plate, and 500ng total plasmids were transfected into the cells. For the CRISPR-ELP plasmid transfection specifically, the molar ratio of each components is dCas9-8xSuntag : gRNA : ScFv-nVenus-ELP : VP64-cVenus-ELP = 2:5:1:1.

### Microscopy sample preparation

For experiments without JF646 staining, cells were seeded onto fibronectin-coated glass-bottom dishes and imaged 4 hours later under a microscope. For experiments with JF646 staining, cells were seeded onto fibronectin-coated glass-bottom dishes with 200nM JF646 containing culture medium. 30 minutes later, cells were gently washed using culture medium and cultured in regular medium or 10μM SAHA containing medium for another 3 hours before imaging. For the heat induced DSB imaging, cells were heated at 43°C for 20min in a thermocycler before JF646 staining and seeding.

### Microscopy, image acquisition, and general processing

Fluorescent live-cell images were taken using the Nikon ECLIPSE-Ti microscope equipped with a 100x, 1.45 numerical aperture (NA) oil immersion objective, a charge-coupled device (CCD) camera, a unique Perfect Focus System (PFS) that automatically corrects focus drift in real-time at 37°C with 5% CO2. For the BFP channel, a 420/40 excitation filter, a 455LP dichroic mirror, and a 480/40 emission filter were used. For the Venus channel, a 495/10 excitation filter, a 510LP dichroic mirror, and a 535/30 emission filter were used. For the HaloTag-JF646 channel, a 630/30 excitation filter, a 660 LP dichroic mirror and a 700/75 emission filter were used. Time-lapse fluorescence images were acquired by MetaMorph 7.8.6.0 software (Molecular Devices, Sunnyvale, California). Z-stack was acquired for all the images. Specifically, for static image, z-stack was acquired for a ±3μm range around the perfect focus at the step of 0.2μm; for the timelapse imaging, to reduce the phototoxicity, z-stack was acquired for a ±3μm range around the perfect focus at the step of 0.5μm. All the z-stack images were projected using the max intensity method in Fiji and subjected to the following analysis.

### Super-resolution imaging (DeepSIM)

DeepSIM super-resolution imaging was performed on a Nikon microscope coupled to the CrestOptics DeepSIM system using a 100× oil-immersion objective (NA 1.49). For each sample, z-stacks were acquired over a 5 μm axial range with a 0.1 μm step size. Images were collected in Deep mode, corresponding to 65 raw images per reconstructed frame. The Venus and JF646 channels were acquired to visualize the CRISPR-ELP droplet and reference signal, respectively.

Raw structured illumination datasets were reconstructed using the DeepSIM software with instrument-matched parameters. Reconstructed image volumes were used to generate 3D renderings and orthogonal xy and xz projections for colocalization assessment.

### Image analysis

Raw image stacks were processed in Fiji/ImageJ. Z-stacks were projected by maximum intensity projection and background subtracted using the rolling ball methods (50 pixels) prior to quantification unless otherwise stated. Detailed analysis is listed below:

### (1) Droplet detection and quantification

Droplet detection and quantification were performed using a custom MATLAB pipeline (MathWorks) based on nucleus segmentation followed by difference-of-Gaussian (DoG) blob detection within nuclear masks. Briefly, single-channel fluorescence images were converted to double precision and normalized to [0,1]. Nuclei were segmented by applying a Gaussian blur (σ = 8 pixels) followed by Otsu thresholding, hole filling, and removal of small objects (minimum nucleus area = 3000 pixels). The resulting nuclear mask was used to restrict puncta detection to nuclear regions.

To detect droplets, images were filtered with two Gaussian kernels (σ_small = 2 pixels; σ_large = 3.5 pixels) and a DoG image was computed (I_DoG = I_σsmall − I_σlarge), producing positive contrast for bright puncta. DoG values within nuclei were used to compute a nucleus-restricted mean (μ) and standard deviation (σ), and candidate puncta were defined as pixels with I_DoG > μ + 3σ. Candidate puncta were further filtered by connected-component analysis and size constraints (area 20–120 pixels²). For each punctum, centroid, area, and mean fluorescence intensity (from the original image) were calculated. Each punctum was assigned to a nucleus by sampling the labeled nuclear mask at the punctum centroid. To quantify droplet signal quality, a local background annulus was defined around each punctum based on its equivalent radius, and background pixels overlapping other detected puncta were excluded. The mean and standard deviation of the local background were then calculated, and punctum signal-to-noise ratio (SNR) was defined as (I_puncta − I_background) / SD_background, where I_puncta is the mean punctum intensity and I_background is the mean intensity of the local annulus. For downstream comparison among conditions, the maximum punctum SNR in each nucleus was used as the cell-level summary metric. For quality control, overlay images showing nuclear boundaries and puncta locations were generated, and per-image as well as combined CSV outputs were exported for downstream analysis.

### (2) Locus enrichment quantification (TetO reference)

To quantify locus-specific enrichment, a 15-pixel diameter circular ROI was centered on the TetR reference punctum and mean Venus intensity was measured. A local background annulus was used to compute normalized enrichment:

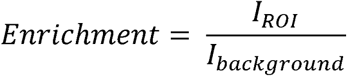

### (3) Particle tracking and mobility analysis

CRISPR-ELP puncta were tracked using TrackMate in Fiji. Mean squared displacement (MSD) and time-averaged MSD (TA-MSD) were computed from tracked trajectories. Apparent diffusion coefficients were estimated from the shortest lag time using:

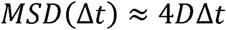

Bound versus unbound puncta were distinguished based on spatial confinement and colocalization with the locus reference.

### (4) 53BP1 dwell time (repair-associated dynamics)

For GLIDRE repair dynamics, 53BP1 recruitment at the locus was quantified by measuring HaloTag-JF646 intensity in an 15×15 pixel rectangle ROI centered on the CRISPR-ELP droplet over time and normalized to local background. Dwell time was defined as the duration during which locus-localized 53BP1 intensity remained 1.5 fold above background from recruitment onset to return.

### RNA extraction and qPCR

U2-OS-SJ cells were transfected with the CRISPR-ELP droplet system, targeting different regions of the target gene. 24 hours later, the cells were FACS sorted based on the Venus signal, and the total mRNA was extracted using the Quick-RNA Microprep Kit (Zymo Research, R1050). Quantitative PCR (qPCR) was performed using iTaq Universal SYBRRTM Green Supermix (Bio-Rad, 1725121) and the mRNA levels were normalized to GAPDH. The primers used in this study are summarized in Supplementary table 3.

### Lentivirus production and stable cell line generation

5×10^6^ HEK 293T cells were seeded in 100cm cell culture dishes one day prior to transfection. A cocktail of pCMV-Δ8.9, pCMV-VSVG, and the transfer plasmid was co-transfected into the HEK 293T cells using the ProFection Mammalian Transfection System (Promega, E1200). 48 hours after transfection, the virus supernatant was collected, filtered through the 0.45μm filter, aliquoted, and frozen in -80°C freezer. To generate the U2-OS-SJ-iCas12a cell lines, 0.25×10^6^ U2-OS cells were seeded into 6-well plate on day before infection. On the second day, 100μl virus supernatant was added into the cells. Five days after infection, cells were heated at 43°C for 20 min, and then sorted based on the mCherry expression level by FACS (SONY SH800).

Primary human T cells were activated using Dynabeads Human T-Expander CD3/CD28 (Gibco, 11141D) at a 1:1 cell-to-bead ratio. At 48 h post-activation, T cells were transduced with lentivirus by spinoculation. Briefly, activated T cells were added together with lentiviral supernatant at a multiplicity of infection (MOI) of 10 to Retronectin-coated plates (Takara, T100A), followed by centrifugation at 1,800 × g for 1 h at 32°C. Dynabeads were removed 5 days after initial activation, and transduced T cells were expanded for an additional 2–5 days before use in experiments or cryopreservation.

### Luciferase-Based Cytotoxicity Assay

Target U2OS cells stably expressing firefly luciferase (Fluc) were seeded at 1 × 10 cells per well in 96-well plates. After 3 h, engineered primary human T cells were added at an effector-to-target (E:T) ratio of 1:3. Co-cultures were incubated for 24 h, after which T-cell-mediated cytotoxicity was quantified by measuring the remaining luminescence of target cells using the ONE-Glo Luciferase Assay System (Promega, E6110), according to the manufacturer’s instructions. Cytotoxicity was calculated from the reduction in luminescence relative to tumor-only control wells.

### Telomere shortening assay

To induce telomere-associated genotoxic stress, H3.3K27WT- and H3.3K27M-expressing U2-OS cells were electroporated with Cas9/sgRNA ribonucleoprotein (Alt-R™ S.p. Cas9 Nuclease V3 and Alt-R™ CRISPR-Cas9 sgRNA, IDT) complexes containing either a telomere-targeting sgRNA or a non-targeting control sgRNA using the Lonza 4D-Nucleofector and the SE Cell Line 4D-Nucleofector X Kit. Following electroporation, cells were plated under standard culture conditions and monitored for 72 h using an IncuCyte live-cell imaging system (Sartorius). Cell growth was quantified from phase-contrast images acquired over time and normalized to the initial time point for each condition. Growth curves were plotted as relative cell growth over 72 h, and the overall proliferative response was summarized by calculating the area under the curve (AUC).

### Flow cytometry

To evaluate heat-inducible AsCas12a expression, U2OS-SJ-iCas12a cells were subjected to heat shock at 43°C for 20 min and returned to 37°C for recovery. At 6 h after heat treatment, cells were harvested and analyzed by flow cytometry. Flow cytometry was performed on a BD Accuri C6 and data were analyzed using FlowJo v10.8.1.

### CAR T spheroid killing assay

U2-OS tumor spheroids were generated using a magnetic field–assisted aggregation approach, following our previously published workflow ^78,79^. Briefly, a magnetic device was assembled with the corresponding μ-Plate 96 Well Square plate (ibidi, Cat. 89621) format to enable rapid and uniform formation of single spheroid per well. Tumor cells were seeded at 20k per well in complete McCoy’s 5A medium (pre-magnetized using 25mM Gadolinium (Bayer)) and placed onto the magnetic device overnight to initiate aggregation. After magnetic assembly, spheroids were maintained for 4 days prior to co-culture.

For co-culture assays, spheroids were incubated with HACVPD-1 CAR T cells at an effector:target (E:T) ratio of 1:1. Co-cultures were imaged every day from Day 1 (1 day after T-cell addition) to Day 3 using microscope. Spheroid size was quantified from GFP images by measuring spheroid area in MATLAB and normalized to Day 1 for each spheroid. For SAHA sensitization experiments, spheroids were pretreated with SAHA (10 μM) for 12 hours and then washed, and then co-cultured with HACVPD-1 CAR T cells as above.

### Public genome track

Publicly available H3K27me3 ChIP-seq, ATAC-seq, SON-TSA-seq, and DamID-seq datasets were used for locus-level epigenomic visualization. DIPG-related H3K27me3 and ATAC-seq bigWig files were obtained from GEO dataset GSE162976. U2OS ATAC-seq bigWig files were obtained from GEO dataset GSE139190. SON-TSA-seq and DamID-seq tracks were obtained from the 4D Nucleome data portal (TSA: 4DNESI8KAJVT; DamID: 4DNESB2HMY7D). Genome tracks from GEO datasets were visualized using the UCSC Genome Browser, whereas 4D Nucleome datasets were visualized using the built-in 4D Nucleome browser. For each locus, relevant bigWig files were loaded as custom tracks and displayed using matched genomic coordinates and scale settings for qualitative comparison of chromatin accessibility, repressive histone marking, and nuclear compartment association.

### Statistical analysis and figure preparation

Statistical analyses were performed using GraphPad Prism. All quantitative measurements were obtained from at least three independent biological replicates unless otherwise stated. For live-cell imaging experiments, individual cells or nuclei were tracked over time and are displayed as data points to illustrate cell-to-cell variability. However, due to the inherently low-throughput nature of long-term single-cell imaging, individual cells derived from the same experiment are not considered fully independent biological replicates. To ensure robustness, all reported trends were reproducible across independent experiments, and statistical analyses were complemented by replicate-level consistency. Normality of data distributions was assessed using the Shapiro– Wilk test. For datasets that did not meet normality assumptions or involved small sample sizes, non-parametric tests (Mann–Whitney or Kruskal–Wallis) were used. Statistical tests were selected as appropriate and are reported in the figure legends. Figures and schematics were assembled using Adobe Illustrator.

## Acknowledgments

We thank Anne Lyons for helpful discussions and for sharing an initial puncta analysis pipeline that informed the early development of our image-analysis workflow. L.Z., and Y.W. designed the research; L.Z., A.M., A.Mo., M.T., X.Y. performed the research. L.Z. and Y.Q. analyzed the data; Z.H., Q.P., and Y.Wu. provided new reagents; L.Z., L.L., and Y.W. wrote the manuscript. All authors reviewed the manuscript and have given approval to the final version of the manuscript. This work was supported in part by grants from NIH R01 CA262815, EB029122, R35 GM140929, R01 HL121365, and HD107206 (Y.Wang.).

## Supporting Information

Supplementary figures S1-10. Supplementary Tables 1-3 and Supplementary movies 1-2 are provided to include related information and resources for this manuscript.

