## Supplementary Figure and Tables for "Programmable Imaging of Locus-Specific DNA Repair Dynamics Reveals Chromatin-Controlled Therapy Resistance in Cancer"

**Supplementary Figures**

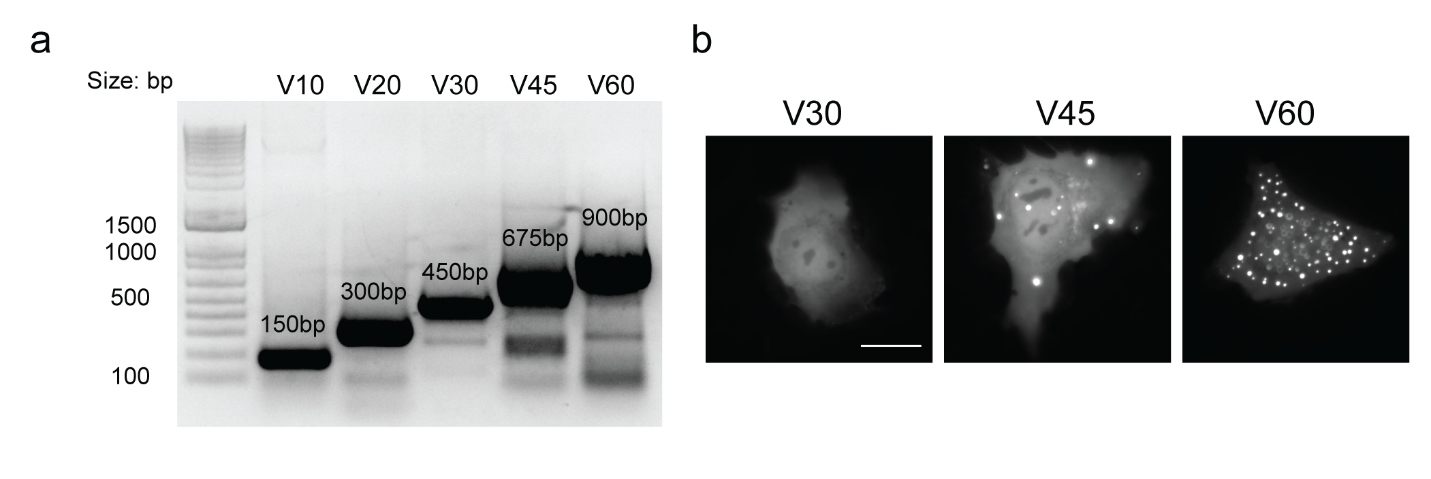

**Supplementary Figure S1. ELP forms droplets in mammalian cells.**

**a.** PCR products showing different sizes of ELP.

**b.** GFP-fused ELP variants display different behaviors in HEK 293T cells. Scale bar: 10μm.

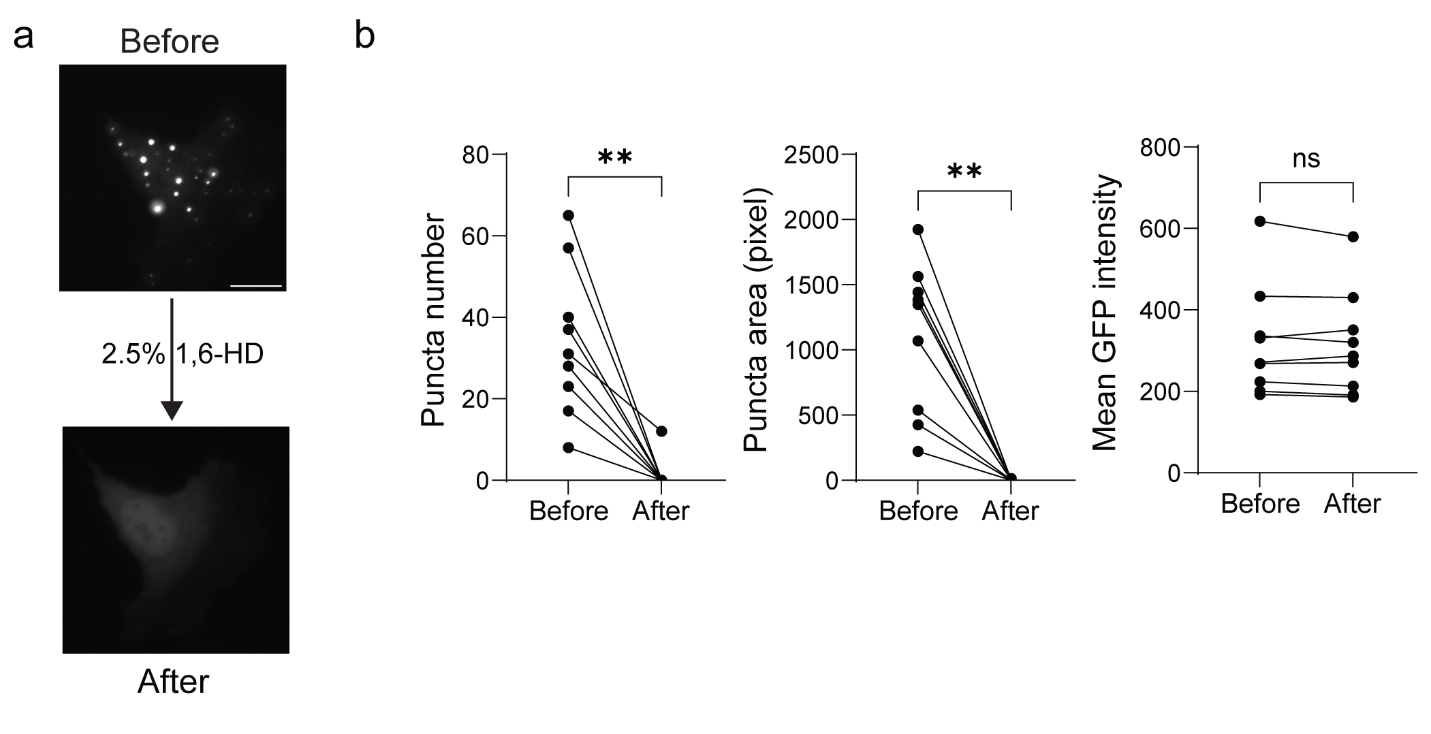

**Supplementary Figure S2. ELP displays the property of liquid-liquid phase separation in mammalian cells.**

**a.** Representative images of GFP-ELP-V60 before and after 2.5% 1,6-HD treatment. Scale bar: 10μm.

**b.** Quantifications of the puncta number, puncta area, and mean GFP intensity before and after 2.5% 1,6-HD treatment. n=9 cells. Wilcoxon test, ** p=0.0039; ns p=0.4258.

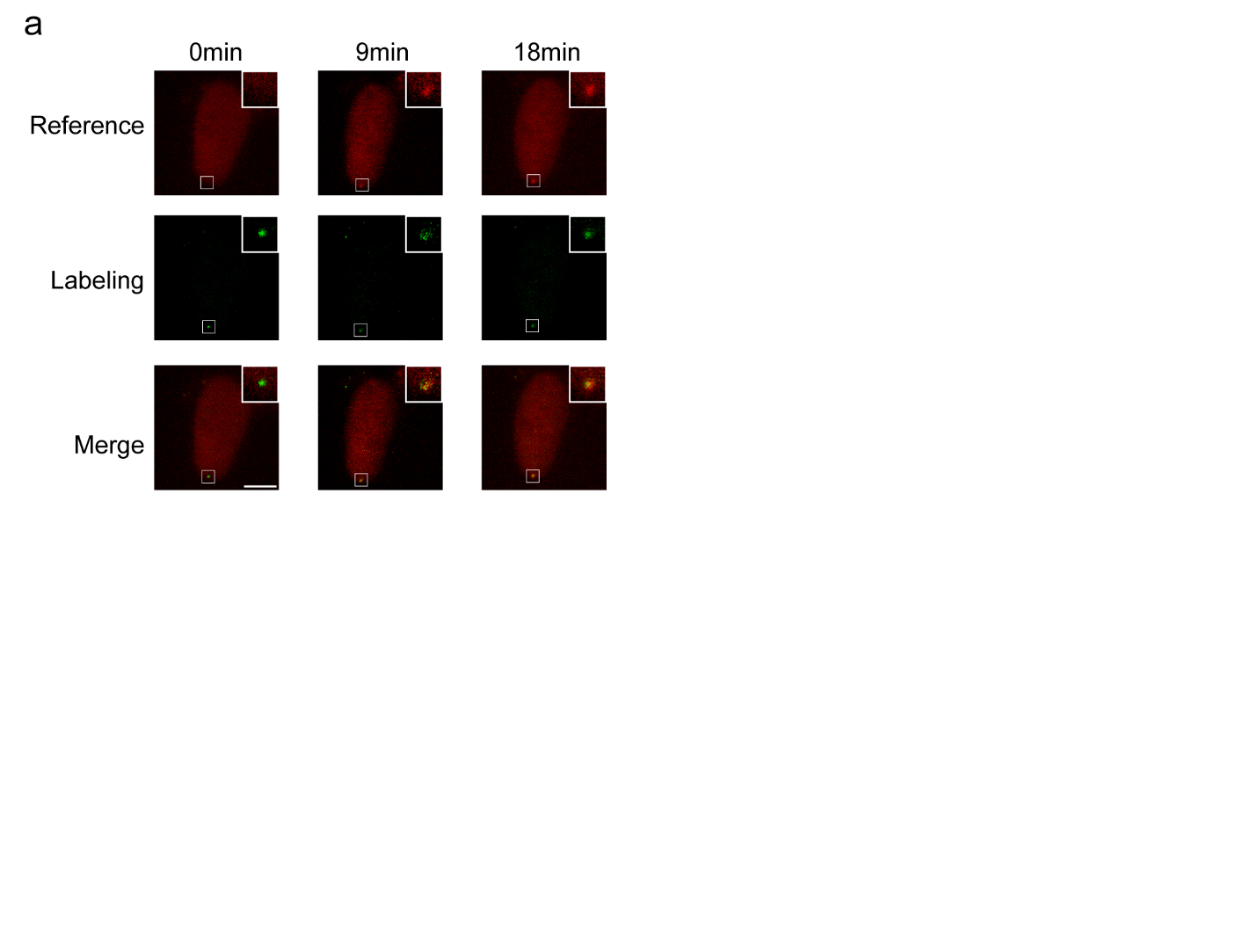

**Supplementary Figure S3. Time-resolved reappearance of TetR-HaloTag signal after doxycycline washout.**

Time-lapse images of U2-OS-SJ cells expressing TetR-HaloTag (reference, red) and CRISPR-ELP droplet labeling signal (green) following doxycycline washout. Images are shown at 0, 9, and 18 minutes after washout, as indicated.

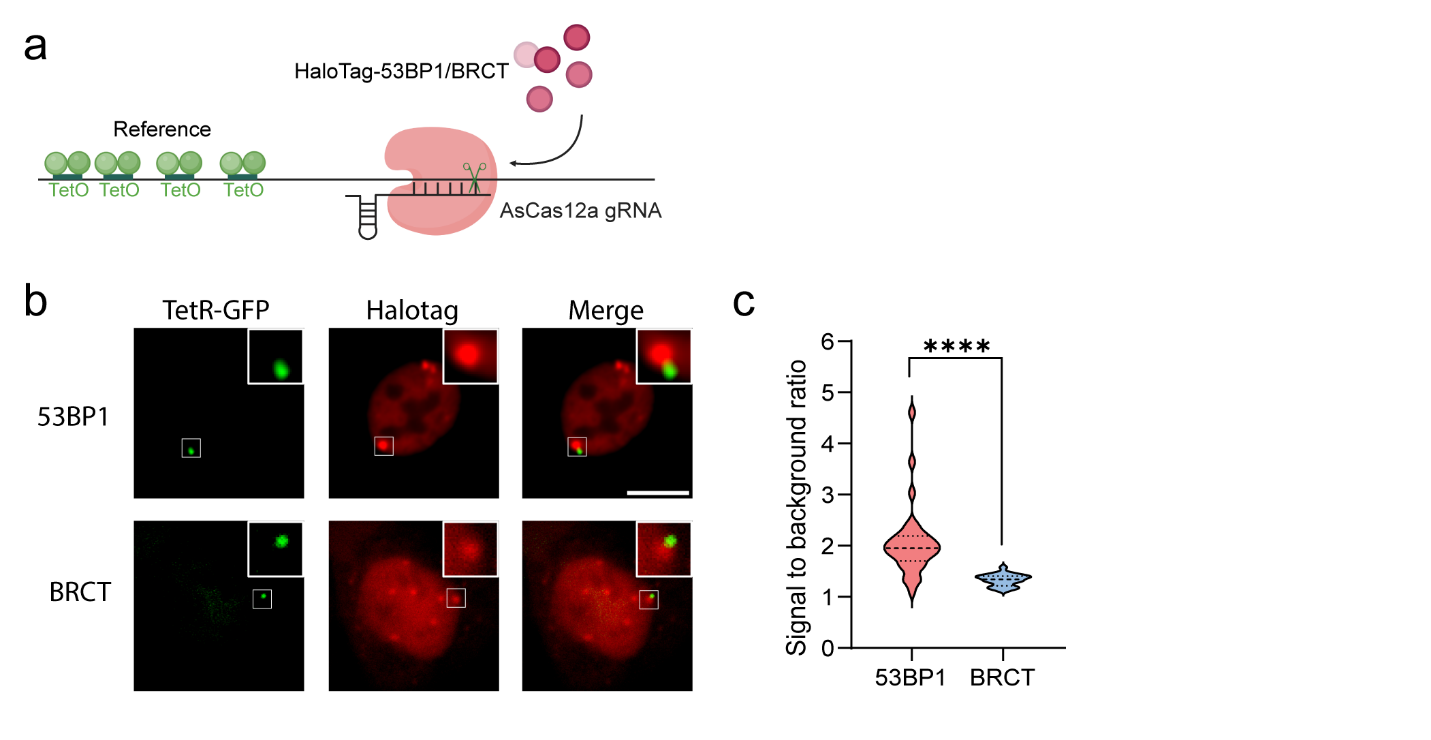

**Supplementary Figure S4. DNA double-strand break can be induced by AsCas12a and monitored by both 53BP1 and the BRCT domain in live cells.**

**a.** AsCas12a and DNA double-strand break monitoring system via 53BP1/BRCT-Halotag. The BRCT domain from the MDC1 protein and the 53BP1 were used to monitor the DNA double-strand break process. The Target was chosen from the chr1p36 region, and TetR-eGFP was used as a reference to indicate the target site.

**b.** Representative images showing 53BP1 and BRCT recruitment at the DSB site. Green, TetR-eGFP reference; Red, 53BP1 or BRCT signal. Scale bar, 10μm.

**c.** Comparison between the 53BP1 and BRCT signal to background ratio at the targeted DSB site. n=33 and 23 respectively, Welch’s t test, **** p<0.0001.

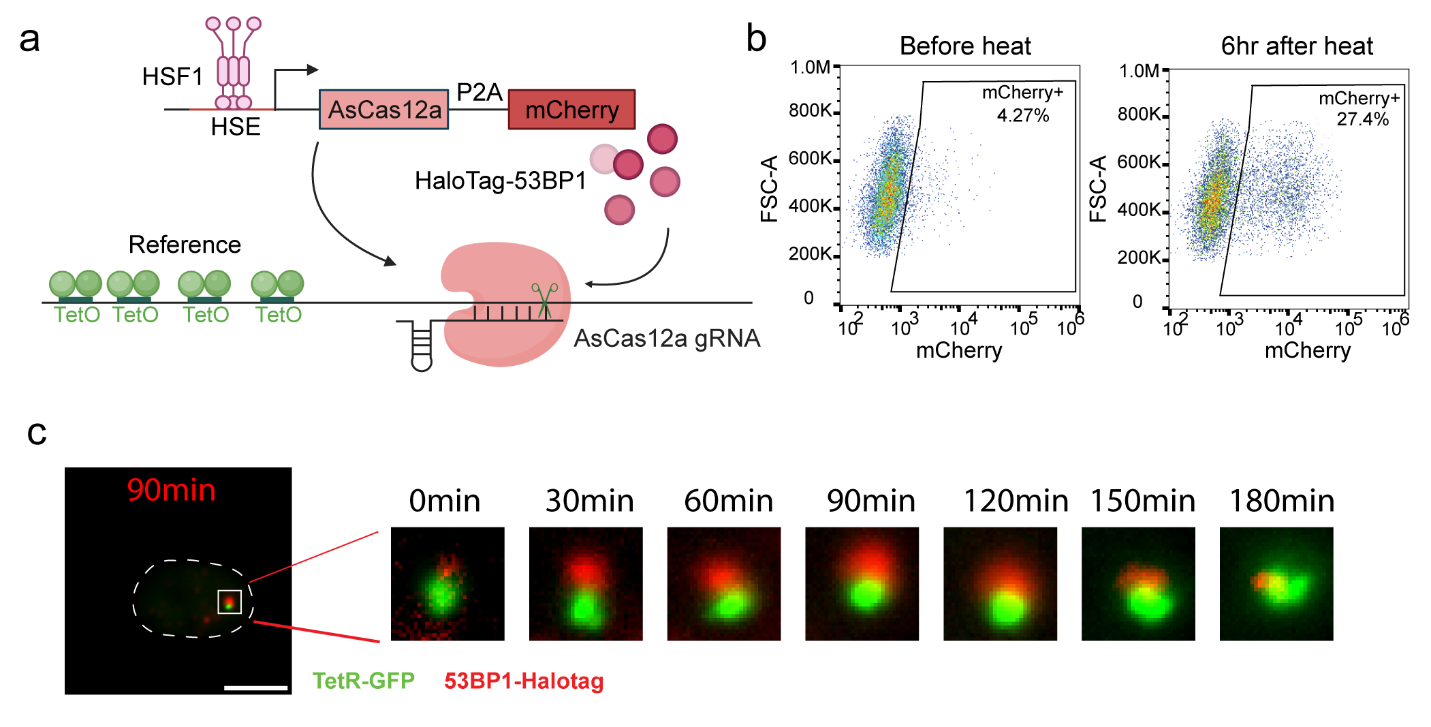

**Supplementary Figure S5. Live cells DSB repair tracking at using TetR-GFP as the reference.**

**a.** AsCas12a is induced by the 7HE-YB promoter upon heat and will be guided to the target non-repetitive locus to induce double-strand DNA break. 53BP1 was used to monitor the DNA double-strand break process. The Target was chosen from the chr1p36 region, and TetR-eGFP was used as a reference to indicate the target site.

**b.** Flow cytometry data showing the inducibility of the AsCas12a indicated by the mCherry signal.

**c.** 53BP1 domain recruitment dynamics. **c**, whole cell image, **d**, enlarged images from the target region. Green, TetR-eGFP reference; Red, 53BP1 domain signal. Scale bar, 10μm.

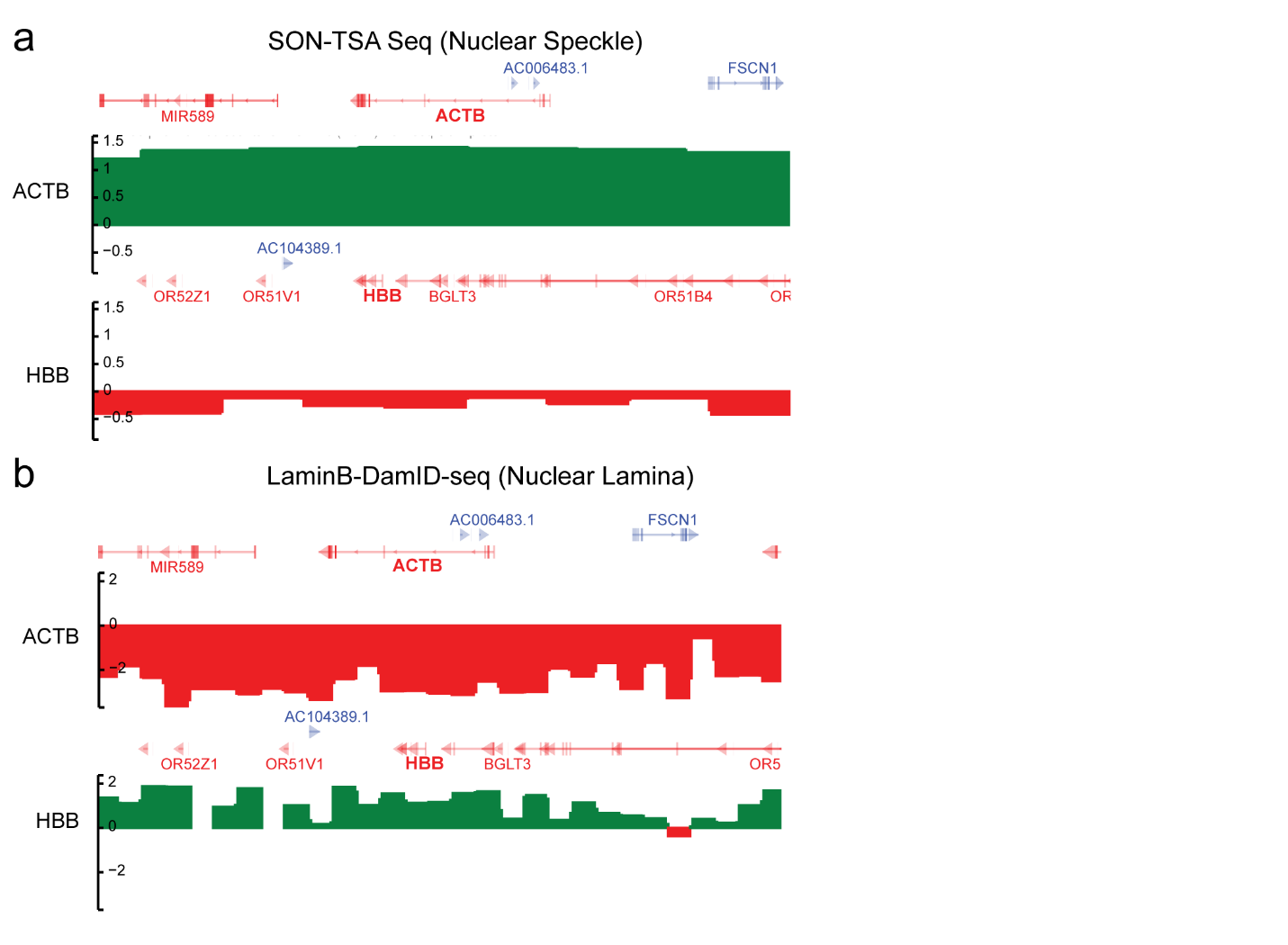

**Supplementary Figure S6. ACTB is nuclear speckle–associated, whereas HBB is nuclear lamina–associated**

**a.** Genome browser snapshots of SON-TSA-seq (nuclear speckle proximity labeling, 4D nucleome database:4DNESI8KAJVT) signal across the ACTB and HBB loci.

**b.** Genome browser snapshots of LaminB-DamID-seq (nuclear lamina association, 4D nucleome database: 4DNESB2HMY7D) signal across the ACTB and HBB loci.

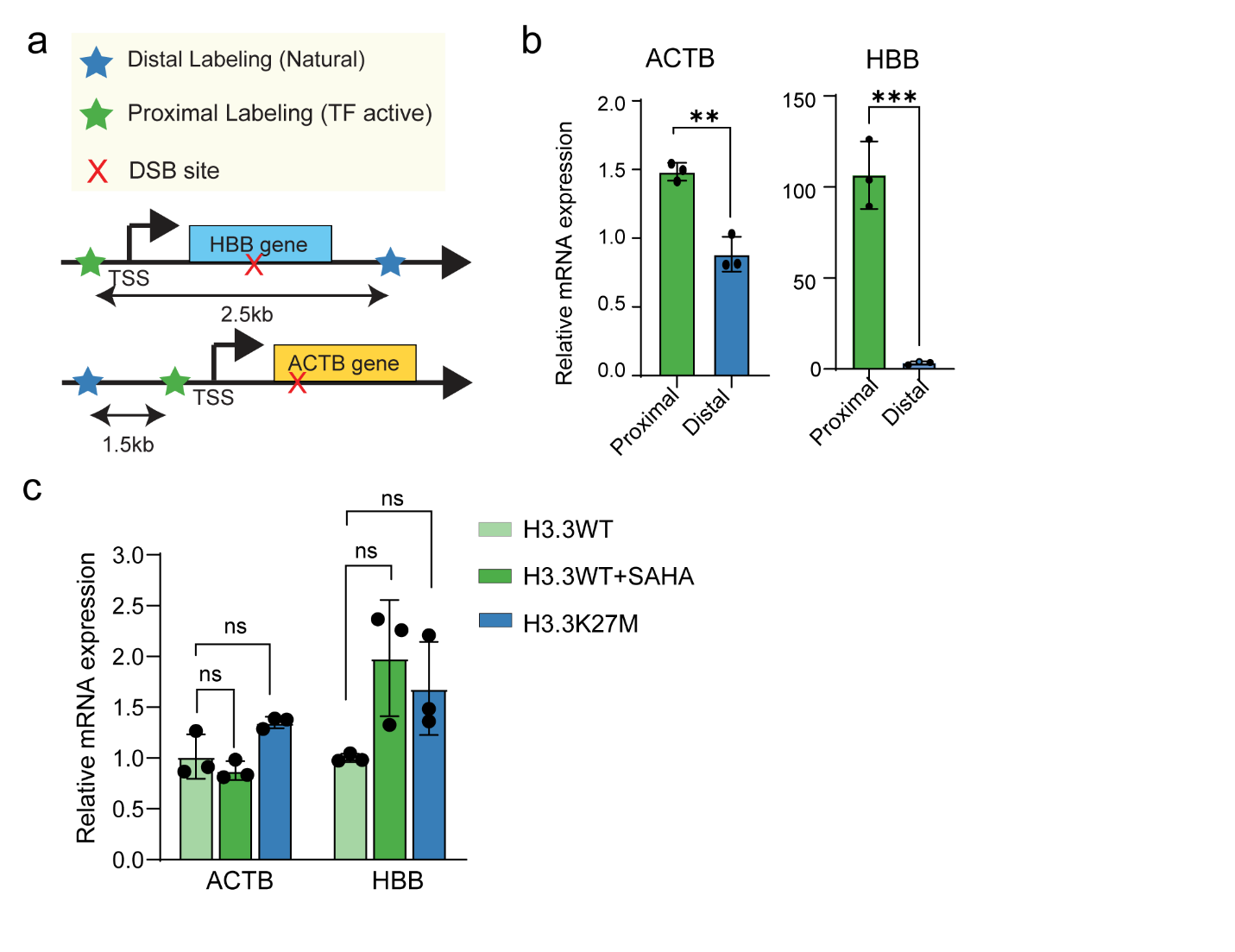

**Supplementary Figure S7. Transcriptional activation and chromatin-opening perturbations at ACTB and HBB**

**a.** Schematic of GLIDRE targeting design for HBB and ACTB. Distal labeling (“Natural,” blue star) and proximal labeling (“TF active,” green star; VP64-functionalized CRISPR-ELP positioned near the TSS) were used to modulate transcriptional output, while the AsCas12a DSB site (red X) was placed within the gene body.

**b.** RT–qPCR quantification of ACTB and HBB mRNA expression under proximal versus distal labeling conditions. Bars show mean ± SD; n=3 technical repeats. Unpaired t-test; ** p=0.0019, *** p=0.0007.

**c.** RT–qPCR quantification of ACTB and HBB mRNA expression under the indicated conditions (H3.3WT, H3.3WT + SAHA, H3.3K27M). Bars show mean ± SD; n=3 technical repeats. Statistical analysis are conducted separately for ACTB and HBB groups. Ordinary one-way ANOVA, ns p>0.05.

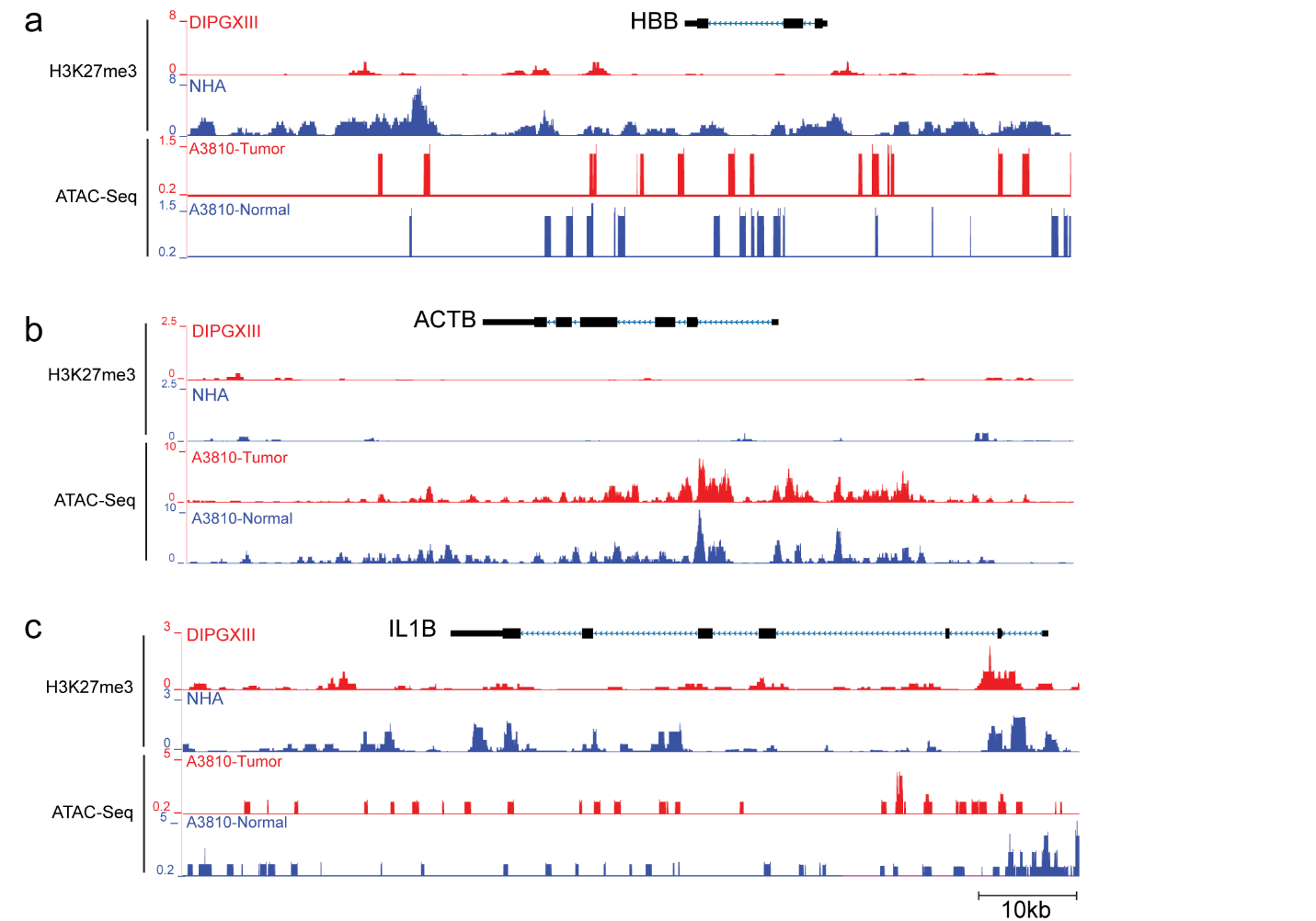

**Supplementary Figure S8. H3K27M-mutant DIPG exhibits locus-dependent changes in chromatin state and accessibility at GLIDRE target genes**

**a-c.** Genome browser snapshots comparing H3K27me3 ChIP-seq and ATAC-seq signals between H3K27M-mutant DIPG cells (red) and normal human cells (blue) at three loci: HBB (a), ACTB (b), and IL1B (c). Gene models are shown above each panel. GSE162976 ^1^.

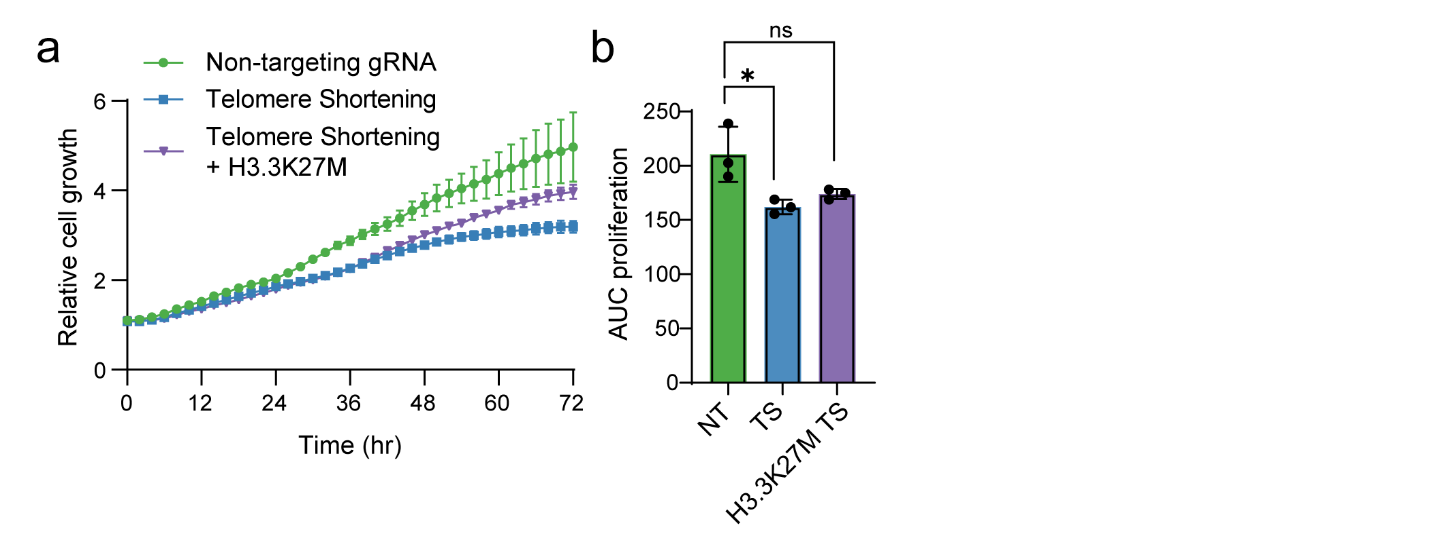

**Supplementary Figure S9. H3.3K27M increases survival under CRISPR–Cas9–induced genotoxic stress.**

**a.** Time-course of relative cell growth over 72 h following CRISPR–Cas9 perturbation normalized to 0hr. Bars show mean ± SEM; n=3 technical repeats.

**b.** Quantification of proliferation by area under the curve (AUC) from the growth curves in (a). NT, non-targeting gRNA; TS, telomere shortening. Bars show mean ± SD; n=3 technical repeats. Kruskal-Wallis test; * p=0.0107, ns p=0.4081.

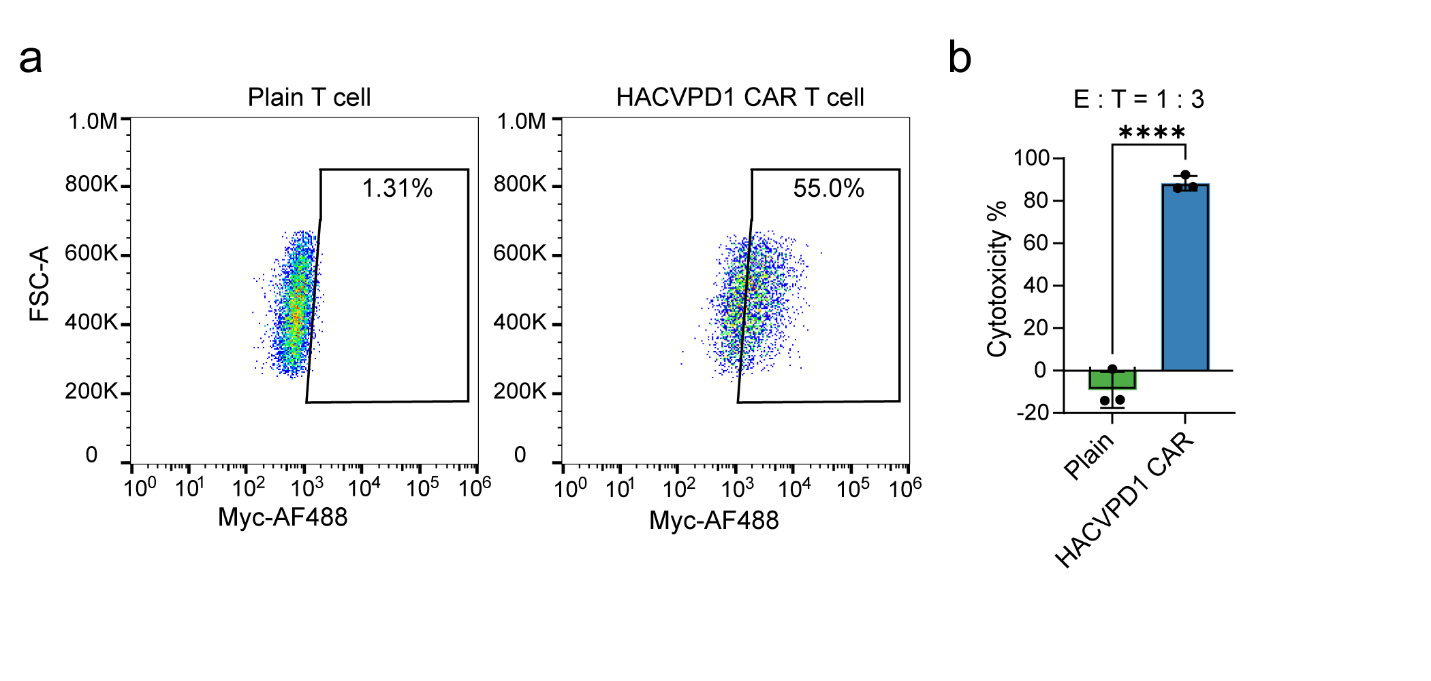

**Supplementary Figure S10. Characterization of HACVPD-1 CAR-T cells and validation of target-cell killing in 2D culture**

**a.** Flow cytometry analysis of HACVPD-1 CAR surface expression on T cells using anti-Myc-AF488 staining. Percentages indicate the fraction of cells within the CAR-positive gate.

**b.** Quantification of cytotoxicity in a 2D luciferase-based killing assay at an effector:target (E:T) ratio of 1:3, comparing plain T cells and HACVPD-1 CAR-T cells against firefly luciferase-expressing U2OS target cells. Cytotoxicity was calculated from the reduction in luciferase signal relative to tumor-only controls. Bars show mean ± SD; n=3 technical repeats. Unpaired Student’s *t*-test; **** p*<*0.0001.

**Supplementary tables**

**Supplementary table 1. List of major plasmids**

| **Plasmid** | **Purpose** | **Source** |
| --- | --- | --- |
| pMAL-c5X-ELP[V-60] | PCR template for ELP variants | Addgene, #67013 |
| dCAS9-ELP-V20 | Single layer labeling | This study |
| pHRdSV40-NLS-dCas9-8x GCN4_v4-NLS-P2A-BFP-dWPRE | The CRISPR-ELP system | This Study |
| pHR-scFv-GCN4-VN155-ELPV20-NLS-dWPRE |  | This study |
| pcDNA-VP64-VC155-ELPV20 |  | This study |
| pHR-scFv-GCN4-eGFP-NLS-dWPRE | Single layer labeling | This study |
| TetR-BFP/GFP/Halotag | TetO site labeling | This study |
| 7HE-YT-AsCas12a-6xNLS-E174R_S542R-p2a-mCherry(pRG232) | Heat-inducible AsCas12a system | This study |
| pCAGG-H3.3K27M | H3.3K27M | This study |
| pcDNA5-FRTTO-Halotag-53BP1 | Tracking DSB | This study |

**Supplementary table 2. List of gRNAs**

| **Target** | **Sequence (5’-3’)** | **Purpose** |
| --- | --- | --- |
| Chr1p36 Single Target | GGAGAGGGGAGGCTGCTGAG | Labeling |
|  | CCCTCAAACTCTCAAGAGCACCT | Cutting |
| IL1B | TGAGATAATTCTCTGGTTCA | Active Labeling |
|  | GTGCAAGTATTTGAGACACG | Inactive Labeling |
|  | TTAGTCCCCTCCCCTAAGAAGCT | Cutting |
| ACTB | AAGCCGGTGAGTGAGCGGCG | Active Labeling |
|  | TCCATCACCACCCCACGCAA | Inactive Labeling |
|  | GCAAAAGGAGGGGAGAGGGGGTA | Cutting |
| HBB | TCAGCCCTCCCTCTAAGATA | Active Labeling |
|  | AACAAAGAAACACTGTGCGA | Inactive Labeling |
|  | GGGATCTGTCCACTCCTGATGCT | Cutting |
| PLK1 | TTGAATAATCAGGACAAAAG | Labeling |
|  | GACTGGCAACCAAAGTCGAATAT | Cutting |
| CDKN2A | GTAAGCACGCGTCACGAGCG | Labeling |
|  | ACTGAGTAGGGCAAATAAATATA | Cutting |

**Supplementary Table 3. List of qPCR primers**

| **Target** | **Sequence (5’-3’)** |
| --- | --- |
| ACTB | CTCTTCCAGCCTTCCTTCCT |
|  | TACAGGTCTTTGCGGATGTC |
| IL1B | CAGAAGTACCTGAGCTCGCC |
|  | AGATTCGTAGCTGGATGCCG |
| GAPDH | AATGGGCAGCCGTTAGGAAA |
|  | GCCCAATACGACCAAATCAGAG |
| HBB | CACCTTTGCCACACTGAGTGAG |
|  | CCACTTTCTGATAGGCAGCCTG |
| PLK1 | GCACAGTGTCAATGCCTCCAAG |
|  | GCCGTACTTGTCCGAATAGTCC |
| CDKN2A | CTCGTGCTGATGCTACTGAGGA |
|  | GGTCGGCGCAGTTGGGCTCC |

**Supplementary Video 1. Heat-inducible AsCas12a enables live-cell visualization of DSB repair-associated dynamics at a genomic locus in U2OS-SJ-iCas12a cells.**

Representative live-cell time-lapse imaging of a target locus in the chr1p36 region marked by TetR-GFP, together with 53BP1-HaloTag as a readout of DSB repair-associated signaling, following heat induction of AsCas12a expression. Cells were heat-shocked at 43°C for 20 min before imaging. Recruitment and resolution of 53BP1 signal at the labeled locus report the onset and progression of the repair response over time.

**Supplementary Video 2. GLIDRE enables live-cell tracking of repair-associated dynamics at a non-repetitive genomic locus.**

Representative live-cell time-lapse imaging of the GLIDRE system at a non-repetitive target locus. CRISPR-ELP droplets label the genomic locus (green), and 53BP1-HaloTag reports repair-associated signaling dynamics (red) following heat-induced AsCas12a cutting.
